# Pharmacological inhibition of nonsense-mediated RNA decay augments HLA class I-mediated presentation of neoepitopes in MSI CRC

**DOI:** 10.1101/2020.10.13.319970

**Authors:** Jonas P. Becker, Dominic Helm, Mandy Rettel, Frank Stein, Alejandro Hernandez-Sanchez, Katharina Urban, Johannes Gebert, Matthias Kloor, Gabriele Neu-Yilik, Magnus von Knebel Doeberitz, Matthias W. Hentze, Andreas E. Kulozik

**Author notes:** To whom correspondence should be addressed (M.W.H.); (A.E.K.).

## Abstract

Microsatellite-unstable (MSI) colorectal cancer is characterized by the accumulation of somatic insertion/deletion (InDel) mutations potentially generating tumor-specific, frameshifted protein sequences. Such mutations typically generate premature translation termination codons targeting the affected mRNAs to degradation by nonsense-mediated RNA decay (NMD), limiting the synthesis and HLA class I-mediated presentation of tumor-specific InDel neoepitopes. We reasoned that the NMD inhibitor 5-azacytidine (5AZA) could serve to increase the expression of NMD-sensitive neoepitopes and analyzed the immunopeptidome of MSI HCT-116 cells using a proteogenomic approach. After immunoprecipitation of HLA:peptide complexes, we identified more than 10,000 HLA class I-presented peptides by LC-MS/MS including five InDel neoepitopes. The InDel neoepitopes were verified on the genomic, transcriptomic, and peptidomic level. Treatment with 5AZA increased the expression of the corresponding frameshift- and premature termination codon-bearing mRNAs and enhanced the presentation of peptides originating from known NMD targets and one of the identified InDel neoepitopes. By analyzing an array of MSI colorectal cancer cell lines and patient samples, we found the underlying frameshift mutation to be highly recurrent and immunization with the corresponding neoepitope induced strong CD8+ T cell responses in an HLA-A*02:01 transgenic mouse model. Our data directly show that peptides originating from frameshifted open reading frames due to InDel mutations in mismatch repair-deficient cells are presented on the cell surface via HLA class I. Moreover, we demonstrate the utility of NMD inhibitor-enhanced HLA class I-mediated presentation of InDel neoepitopes as well as their immunogenicity, uncovering the clinical potential of NMD inhibition in anti-cancer immunotherapy strategies.

**One Sentence Summary:** Immunopeptidomics identified increased HLA class I-mediated presentation of immunogenic, frameshift-derived neoepitopes following NMD inhibition.

## Introduction

Microsatellite-unstable colorectal cancers (MSI CRCs) account for approximately 15% of all CRCs, and hence about 275.000 cases per year (*1*). These cancers are caused by inactivation of the DNA mismatch repair (MMR) system, either by sporadic epigenetic silencing of *MLH1* or by the combination of inherited, monoallelic germline mutations and a second hit in MMR genes (a cancer predisposition termed Lynch syndrome)(*2*). As a consequence, MSI CRCs are characterized by the accumulation of somatic mutations, mainly small insertion/deletion (InDel) mutations in repetitive DNA stretches termed microsatellites (*3*). InDel mutations in protein-coding microsatellites typically lead to functional inactivation of the affected genes and promote tumorigenesis if suppressor genes are disrupted (*4*). Importantly, two-thirds of all InDel mutations cause a shift of the reading frame, hence encoding tumor-specific protein sequences (*2*). Endogenously synthesized proteins that pass through the cellular antigen processing and presentation machinery are ultimately presented at the surface of all nucleated cells by human leukocyte antigen (HLA) class I molecules. HLA class I molecules bind peptides with a length of eight to 15 amino acids (AAs) via allele-specific anchor positions at the peptides’ C- and N-termini, while exposing the central part to cells of the immune system. HLA class I-presented peptides (HLAp) derived from frameshifted protein sequences encoded by microsatellites carrying InDel mutations (InDel neoepitopes) allow patrolling CD8^+^ T cells to identify and target cells presenting such neoepitopes (*5*). In contrast to neoepitopes derived from a single AA change (SNP neoepitopes), frameshifted, tumor-specific protein sequences can potentially harbor several InDel neoepitopes with binding capacity to different HLA allotypes and InDel neoepitopes have been suggested to possess higher immunogenicity caused by their fundamental difference to endogenous self-antigens originating from wild-type proteins (*6*). Several studies support the presence of InDel neoepitope-specific cytotoxic T cells in both, healthy individuals and MSI CRC patients (*7, 8*).

Recently, we showed in a clinical trial the induction of neoepitope-directed immune responses after vaccination with shared, *in silico* predicted neoepitopes in patients with MMR deficiency (*9*). Moreover, it was shown that T cells, re-activated by immune checkpoint inhibition, target tumor-specific neoepitopes thus further emphasizing the crucial role of neoepitopes in immunotherapeutic strategies (*10*). However, both strategies rely on the expression and presentation of such neoepitopes in sufficient quantities. Cancer cells constantly evolve to evade the immune system by various mechanisms such as decreasing the expression of HLA class I molecules, increasing the secretion of inhibitory cytokines and ligands, or by reducing the quantity and quality of presented antigens (*11*).

The expression of most InDel neoepitopes is limited by nonsense-mediated RNA decay (NMD). NMD is a conserved quality control pathway that recognizes and degrades mRNAs with premature termination codons (PTCs) introduced by nonsense or frameshift mutations, transcription errors, incorrect splicing, or unprogrammed ribosomal frameshifting. NMD thus limits the synthesis of C-terminally truncated and potentially harmful proteins (*12*). However, NMD also regulates the expression of error-free, physiologic mRNAs (so-called endogenous NMD targets) with characteristic NMD-stimulating features such as upstream open reading frames, exceptionally long 3’UTRs, splice junctions in the 3’UTR, or programmed ribosomal frameshifting (*13*). Such endogenous NMD targets contribute to key biological processes such as embryogenesis, organ development, and stress responses (*14*). In MSI CRCs, the central NMD factors *UPF1*, *UPF2*, *SMG1*, *SMG6,* and *SMG7* are expressed substantially more strongly compared to microsatellite-stable CRCs and it has been suggested that increased NMD activity can promote MSI tumorigenesis by degrading mutated transcripts (*15, 16*). Because InDel mutations often introduce NMD-triggering PTCs in the frameshifted, alternative reading frames, NMD restricts the production and consequently the immune recognition of InDel neoepitopes (*15–17*). It is known that both, pharmacological and siRNA-mediated NMD inhibition stabilize InDel-mutated transcripts in MSI CRC (*15, 17*). Furthermore, NMD efficiency negatively correlates with host immunity against MSI CRC (*15, 16*).

Leveraging the potential of InDel neoepitopes in generating effective T cell responses for novel and eventually personalized immunotherapy strategies requires the reliable identification of these peptides. Recent breakthroughs in the sensitivity and reproducibility of mass spectrometry (MS) enable the unbiased exploration of the global immunopeptidome presented by the HLA system. This novel methodology also enables the identification of HLA-presented neoepitopes and helps to overcome disadvantages of previous methods that were based on HLA binding predictions and indirect immunological read-outs (*18*). New methodological MS approaches such as dual fragmentation by electron-transfer/higher-energy collision dissociation (EThcD) both expand the detectable immunopeptidome and increase the confidence in peptide identifications (*19*). Furthermore, *de novo* peptide sequencing allows the identification of neoepitope sequences not included in standard proteomics databases (*20*). However, most InDel neoepitopes are NMD targets, which limits their abundance and thus their recognition by the immune system. Recently, our laboratory identified 5-azacytidine (5AZA) as a potent NMD inhibitor (*21*). Importantly, 5AZA inhibits NMD without interfering with protein synthesis at therapeutic concentrations distinguishing 5AZA from other known NMD inhibitors (*21*). Therefore, therapeutic NMD inhibition by 5AZA could, in theory, help to increase the production of InDel neoepitopes and increase tumor recognition by the host’s immune system. Here, we report the first unbiased, direct identification of previously unknown, immunogenic InDel neoepitopes in MSI colorectal cancer by mass spectrometry, and provide experimental evidence that NMD inhibition increases the HLA class I-mediated cell surface presentation of InDel neoepitopes.

## Results

### Validation of the experimental system

The MSI CRC cell line HCT-116 was chosen as the starting model system for this study. HCT-116 cells express six different HLA class I alleles, including the common HLA-A*02:01 allele, allowing the presentation of a broad spectrum of peptides (*22*). NMD competence of HCT-116 cells was determined using a transiently transfected dual-luciferase reporter system (*23*). HCT-116 cells exhibit a high NMD efficiency demonstrated by the substantially and highly significantly lower *Renilla* luciferase signal in cells transfected with the NS39 reporter (0.055 ± 0.038 normalized to WT reporter signal, *p* ≤ 0.0001)(Supplementary figure 1). The NMD restricting effect of 5AZA in HCT-116 cells was tested by assessment of transcript levels of known endogenous NMD targets by quantitative real-time PCR (qPCR). Treatment with 5 µM 5AZA for 24 h induced a significant stabilization (*p* ≤ 0.0001) of ATF3 (3.6-fold), SC35C (2.9-fold), and SC35D (2.7-fold) transcripts (Supplementary figure 1).

Conventionally, neoepitopes are identified by evaluating *in silico* predicted candidates using T cell screening technologies. However, such analyses are limited by high rates of false-positive results (*24–26*). By contrast, MS-based immunopeptidomics offers the only unbiased method to directly identify (neo-)epitopes that are actually presented via HLA class I molecules on cancer cells. We extended a recently published, high-throughput workflow for the identification of HLA-presented peptides (*27*) to enable the identification of InDel neoepitopes and investigate the effect of NMD inhibition at the level of the immunopeptidome (Figure 1). Briefly, after IP with a pan-HLA antibody (Supplementary figure 2) and subsequent separation of the bound peptides from HLA class I molecules, HLAp were analyzed by LC-MS/MS applying different fragmentation methods. Higher-energy collisional dissociation (HCD) is the standard fragmentation mode for acquiring high-resolution data at a fast speed and therefore provides in-depth coverage of the immunopeptidome. Electron-transfer/higher-energy collision dissociation (EThcD), a combination of HCD and electron transfer dissociation (ETD), generates more complex fragmentation spectra leading to a higher peptide sequence coverage to ensure high-confidence identification. Furthermore, we combined fragmentation by EThcD with precursor selection targeting low abundance precursors first (lowEThcD) to compensate for the lower coverage caused by the slower acquisition frequency of EThcD fragmentation. Finally, the obtained MS raw data were subjected to a *de novo* sequencing-assisted database search which improves both, sensitivity and accuracy of peptide identifications and enables the identification of neoepitopes.

**Figure 1:**
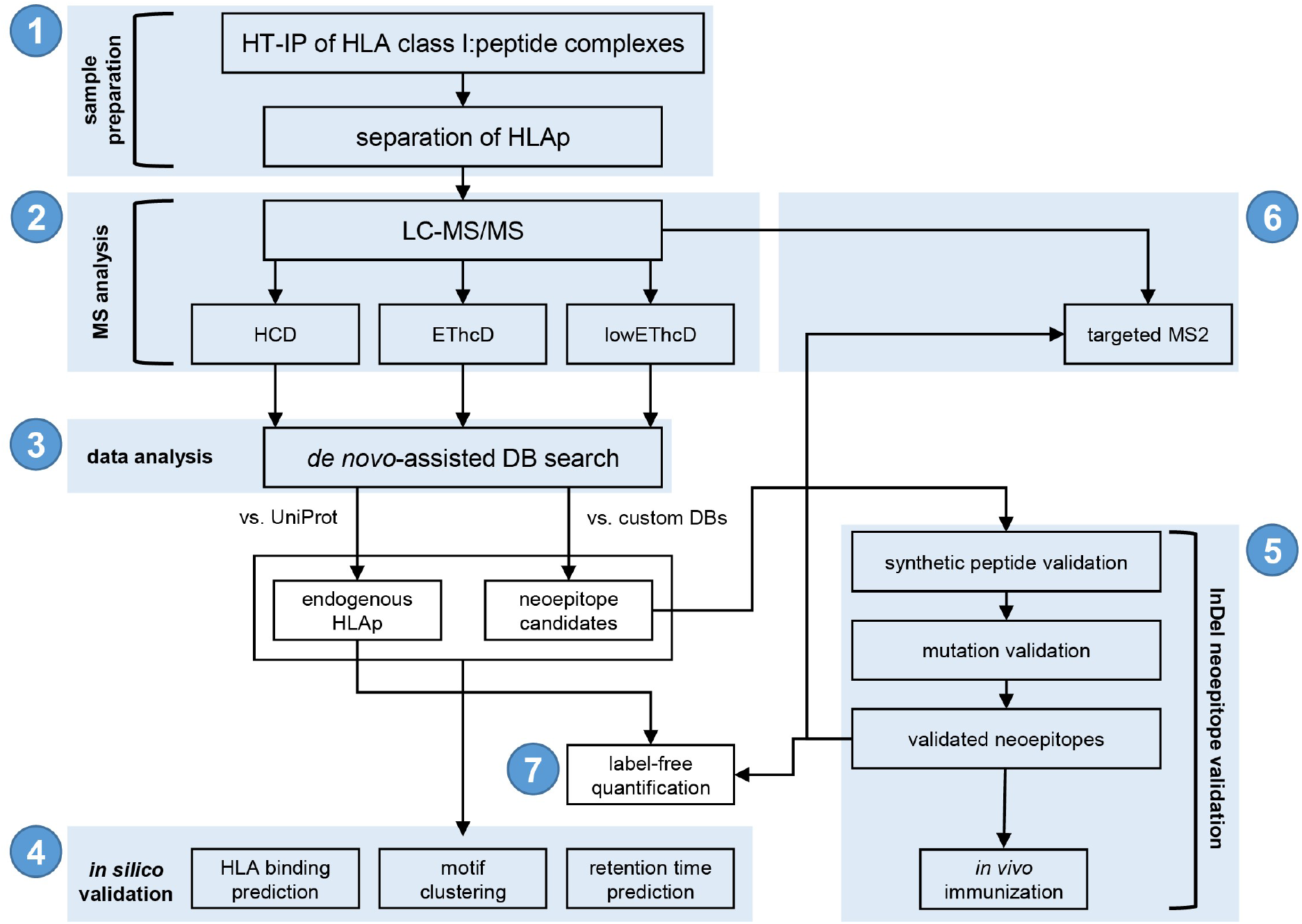
Immunopeptidomics workflow for the identification, validation, and quantification of InDel neoepitopes. After high-throughput immunoprecipitation of HLA class I:peptide complexes, HLAp are separated (1) and subjected to LC-MS/MS using different fragmentation and precursor selection methodologies (HCD, EThcD, lowEThcD; 2). Data analysis of raw data is performed using de novo-assisted database search against the UniProt database to identify endogenous HLAp and against custom databases containing neoepitope sequences (3). All identified peptides are validated bioinformatically (binding prediction, sequence clustering, retention time prediction; 4). Neoepitope candidates are further validated by comparison to synthetic peptide spectra and validation of underlying mutations. *In vivo* processing and presentation are tested by immunization of an “HLA-humanized” mouse model (5). Neoepitopes are measured again using a targeted MS2 approach (6). Label-free quantification is performed for endogenous HLAp and validated neoepitopes (7). HT-IP, high-throughput immunoprecipitation; HLAp, HLA class I-presented peptides; MS, mass spectrometry; LC-MS/MS, liquid chromatography-tandem mass spectrometry; DB, database.

### Direct, mass spectrometry-based identification of HLA class I-presented peptides

The workflow described above was used to analyze HLAp isolated from HCT-116 cells treated either with 5 µM 5AZA or the solvent DMSO control for 24 h. We identified a total of 10,030 unique HLAp at a stringent false discovery rate (FDR) of 1% (Figure 2A). Of these, 3,098 HLAp (31% of the total dataset) were identified both in datasets recorded using HCD or dual fragmentation (EThcD/lowEThcD), while 4,907 HLAp (49%) were only identified using HCD fragmentation. 2,025 HLAp (20%) were only identified in the dual-fragmentation datasets. MS1 signals of identified and quantified peptides show a wide range of intensity spanning several orders of magnitude (log2 intensity: 12.75 – 32.55, mean log2 intensity: 20.19). Peptides identified by all three fragmentation methods show a slightly higher average intensity/abundance when compared to peptides identified by HCD fragmentation alone (log2 intensity: 21.58 vs. 19.77). Of note, we identified a subset of 851 low abundance peptides (mean log2 intensity: 18.52) using the lowEThcD method, which preferentially targets less abundant peptide precursors (Figure 2A). In summary, these findings illustrate the benefit of applying both different fragmentation methods and precursor selection strategies to increase the number of peptide identifications providing a more comprehensive view of the immunopeptidome. To ensure the quality of the obtained dataset, various *in silico* quality controls were performed. First, we calculated the sequence-specific hydrophobicity index (HIs), which is an orthogonal parameter to validate correct peptide identifications and correlates with experimentally observed retention times (*28*). HIs of identified peptides showed a tight correlation with observed retention times for all three fragmentation methods used (Pearson’s correlation coefficient HCD: 0.96, EThcD: 0.96, lowEThcD: 0.95)(Figure 2B). Next, we analyzed the HLA-associated properties of the identified peptides. The peptides showed an HLA class I-typical length distribution with mainly nonamers (Figure 2C). Investigation of the entire immunopeptidome using MS reduces the *a priori* introduced bias of selectively surveying (neo)epitopes shortlisted by *in silico* binding predictions. However, binding prediction of identified HLAp *a posteriori* represents a suitable validation tool for immunopeptidomics datasets. Using NetMHCpan, we found that 90% of the identified peptides (8,988 peptides; 7,286 SB, 1702 WB) were predicted to bind at least one of the HLA class I molecules expressed on the HCT-116 cells (Figure 2D). Of the predicted binders, 21% showed the highest affinity for HLA-A*01:01 (1,912 peptides; 1,816 SB, 96 WB), 18% for HLA-A*02:01 (1,590 peptides; 1,098 SB, 492 WB), 19% for HLA-B*18:01 (1,742 peptides; 1,326 SB, 417 WB), 31% for HLA-B*45:01 (2,788 peptides; 2,336 SB, 452 WB), 9% for HLA-C*05:01 (767 peptides; 582 SB, 185 WB) and 2% for HLA-C*07:01 (188 peptides; 128 SB, 60 WB). Considering the similarity of HLAp which is caused by their allele-specific binding-mediating anchor residues, we clustered peptide sequences into groups and identified four distinct motifs that correspond to the consensus binding motifs of HLA-A*01:01, HLA-A*02:01, HLA-B*18:01, and HLA-B*45:01 (Figure 2E). Although expressed on HCT-116 cells, consensus binding motifs for HLA-C*05:01 and HLA-C*07:01 could not be defined probably due to the low cell surface expression of the corresponding alleles and their motif redundancy to HLA-A and B alleles (*26, 29*). Finally, we analyzed the source proteins of the identified peptides. Of the 4,767 distinct source proteins, 2,524 (53%) were represented by only one, 1,054 (22%) by two, and 516 (11%) by three distinct HLAp at the cell surface. The remaining 673 source proteins (14%) were represented by four or more distinct HLAp (Supplementary figure 3). In general, source proteins were associated with a broad spectrum of cellular localization. In line with previous reports, the source proteins of the top 10% most abundant HLAp were significantly enriched in clusters for nuclear, cytoskeletal, and ribosomal proteins (*26, 30*). Taken together, these data validate our dataset as a representative view of the endogenous immunopeptidome of HCT-116 cells.

**Figure 2:**
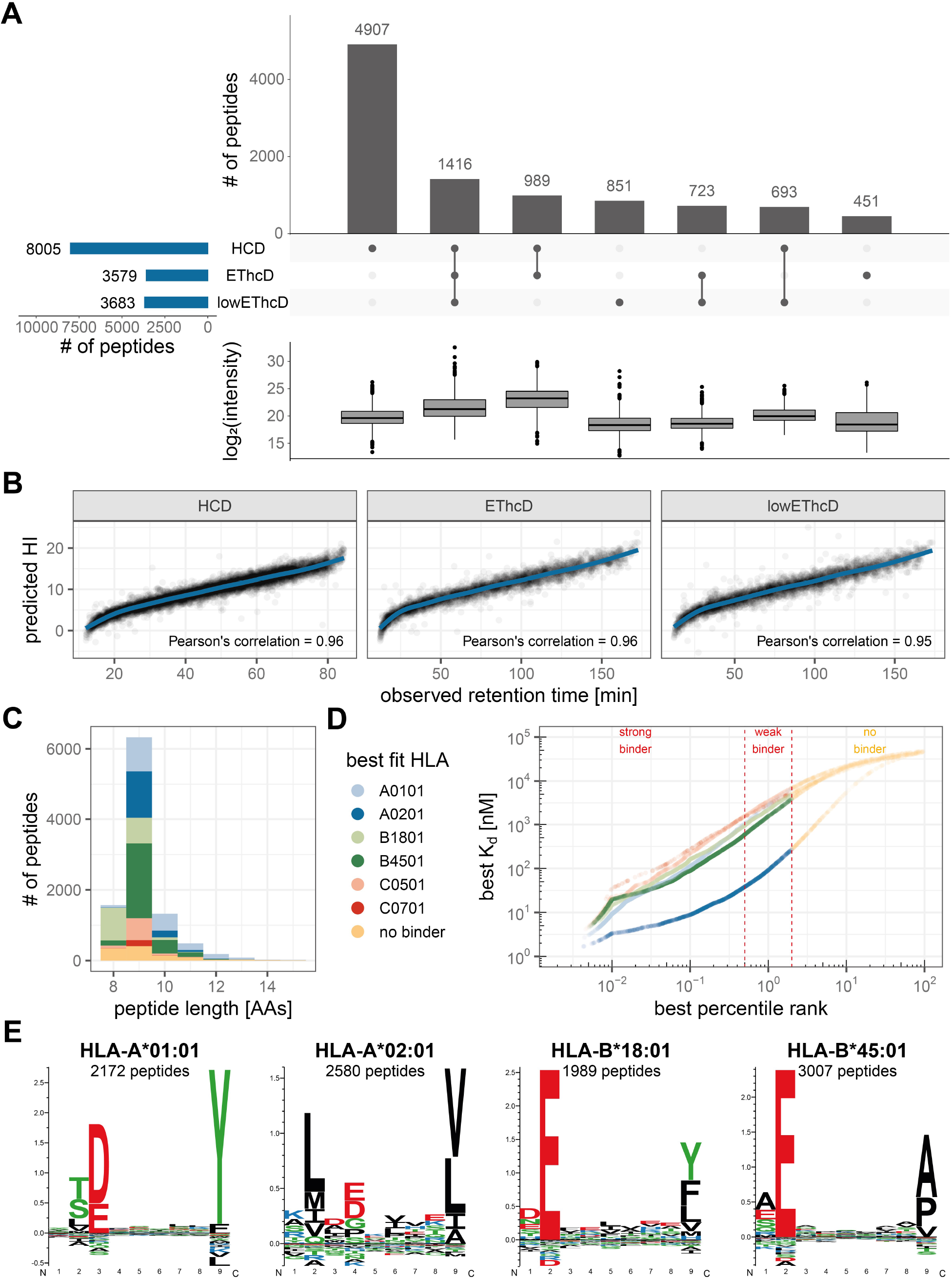
Quality control and characteristics of identified peptides. **(A)** Number of peptides identified using different MS fragmentation methods and overlapping sets. Boxplot summary representing intensity distribution for subsets of peptides. **(B)** Predicted hydrophobicity index (HI) against observed retention time of identified peptides for different MS fragmentation methods. **(C)** Typical length distribution of HLA class I-presented peptides. Colors represent best fitting HLA allele determined by NetMHCpan 4.0. **(D)** Binding prediction of all identified peptides. Threshold for strong binders is top 0.5% ranked, for weak binders top 2%. **(E)** Sequence clustering of identified peptides to four distinct binding motifs matching HLA alleles expressed by HCT-116 cell line. 282 outliers (2.8%) were not clustered.

### Identification and validation of HLA class I-presented InDel and SNP neoepitopes

After having created a high-quality, representative dataset of endogenous HLAp, we next sought to query this dataset for the existence of HLA class I-presented neoepitopes. To enable the identification of both InDel and SNP neoepitopes we constructed custom, cell line-specific databases based on publicly available sequencing data. InDel databases based on COSMIC and CCLE mutation data for the HCT-116 cell line contain 883 unique entries. The lengths of frameshifted, tumor-specific protein sequences range from 13 to 393 AAs (median: 31 AAs), potentially generating 26,298 unique peptides with a length of nine AAs, which is the preferred binding length of HLA class I molecules. Of these, 11% (2,782 peptides; 911 SB, 1,871 WB) were found to potentially bind at least one of the HLA alleles expressed by HCT-116 cells (Supplementary figure 4). The SNP database contains 1,260 previously reported *in silico* predicted potential SNP neoepitopes (*22*).

In contrast to standard MS data analysis workflows, PEAKS reports spectra with high *de novo* sequencing scores, which were not matched to a UniProt database entry as “*de novo* only” spectra. These high-scoring “*de novo* only” spectra were subsequently searched against the custom SNP and InDel databases. After applying a stringent FDR of 1%, we identified nine InDel-and five SNP-neoepitope candidates (Figure 3A). Three of these five SNP neoepitopes have been reported previously (*26*). To validate the identifications of the neoepitope candidates, we first used BLASTp with standard parameters for short input sequences to rule out that identified peptides (and leucine/isoleucine permutations of them) match known human AA sequences. We excluded one of the nine InDel neoepitope candidates matching the 14 AAs of the wild-type protein part which was included during database generation (Figure 3A).

**Figure 3:**
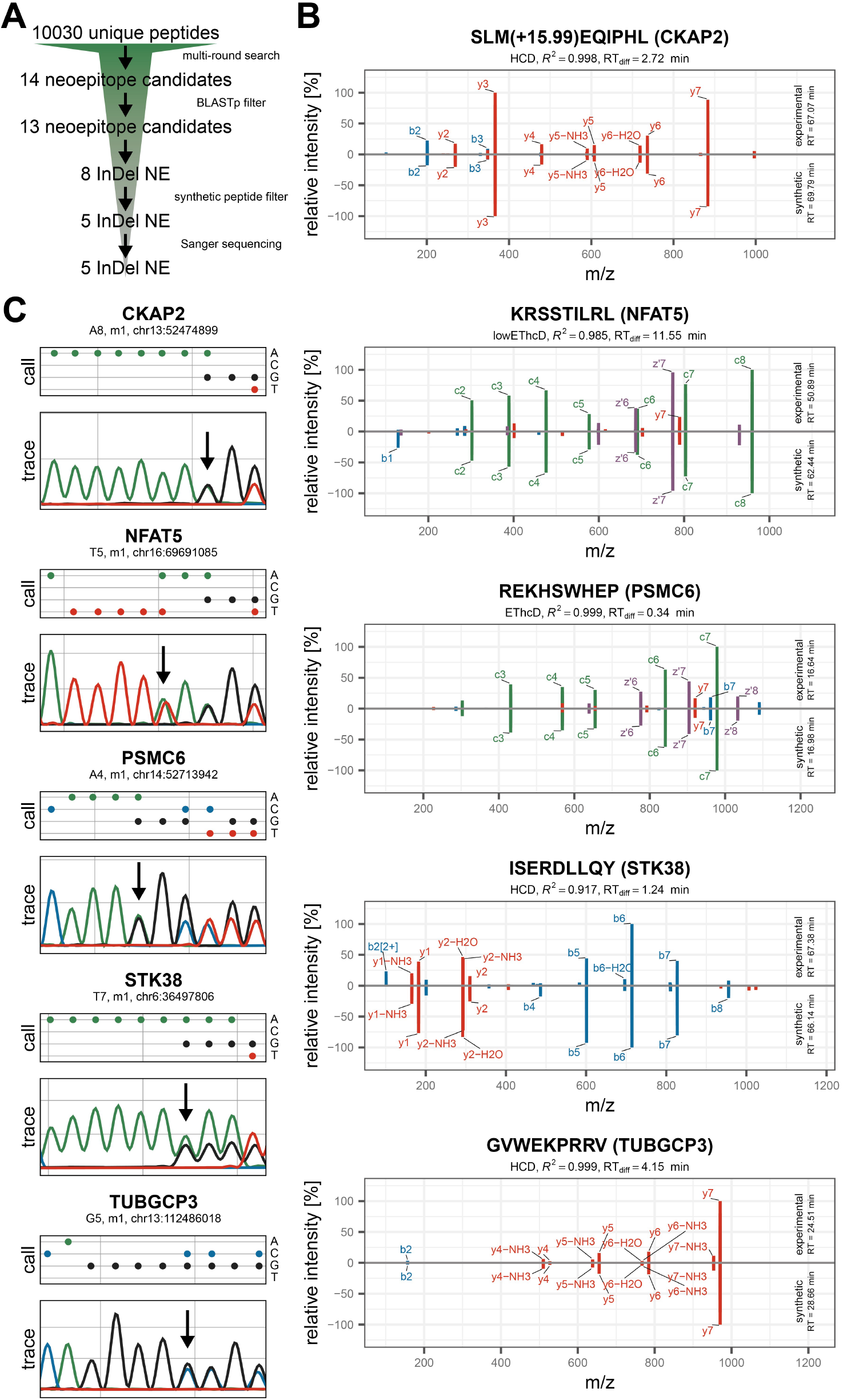
Validation of identified InDel neoepitopes. **(A)** Overview of validation procedure. Candidates were filtered using BLASTp to exclude peptides matching endogenous proteins. Spectra of candidates were compared to spectra recorded from synthetic peptides and underlying frameshift mutations were confirmed by Sanger sequencing. **(B)** Comparison of matched ions observed in candidate spectra (top) and synthetic peptide spectra (bottom). Top 10 most intense ions are labeled, retention time difference and correlation between experimental and synthetic peptide spectrum is reported. **(C)** Base calls and sanger traces of underlying frameshift mutations. Positions of InDel mutations are indicated by an arrow. m1, minus one base pair deletion.

To confirm the identity/AA sequence of neoepitopes, we next compared their spectra with those obtained from synthetic peptide counterparts. The MS acquisition and data analysis workflow previously used for the identification of neoepitopes from cell line samples was applied to pools of synthetic peptides and confirmed the identities of five InDel neoepitopes and five SNP neoepitopes (Figure 3B/Supplementary figure 5B). Intensities of matched fragment ions showed very high correlations (Pearson’s correlation: 0.917 – 0.999) for correct identifications using both, HCD and EThcD fragmentation methods, while this correlation was much lower for the four false-positive identifications (Pearson’s correlation: 0.073 – 0.673). Furthermore, we observed substantial differences in retention times between experimental samples and synthetic peptide pools for the false-positive identifications. Binding prediction for identified InDel and SNP epitopes showed that all but one of the validated peptides are predicted to bind at least one of the HLA alleles expressed on HCT-116 cells. Furthermore, we confirmed the underlying mutations for all validated InDel and SNP neoepitopes in the genomic DNA using Sanger sequencing (Figure 3C/Supplementary figure 5C). Taken together, these data show the validity of the identification of both InDel and SNP neoepitopes. Table 1 and 2 provide an overview of all validated HLA class I-presented InDel and SNP neoepitopes and the biological function of the InDel neoepitope source proteins.

**Table 1:**
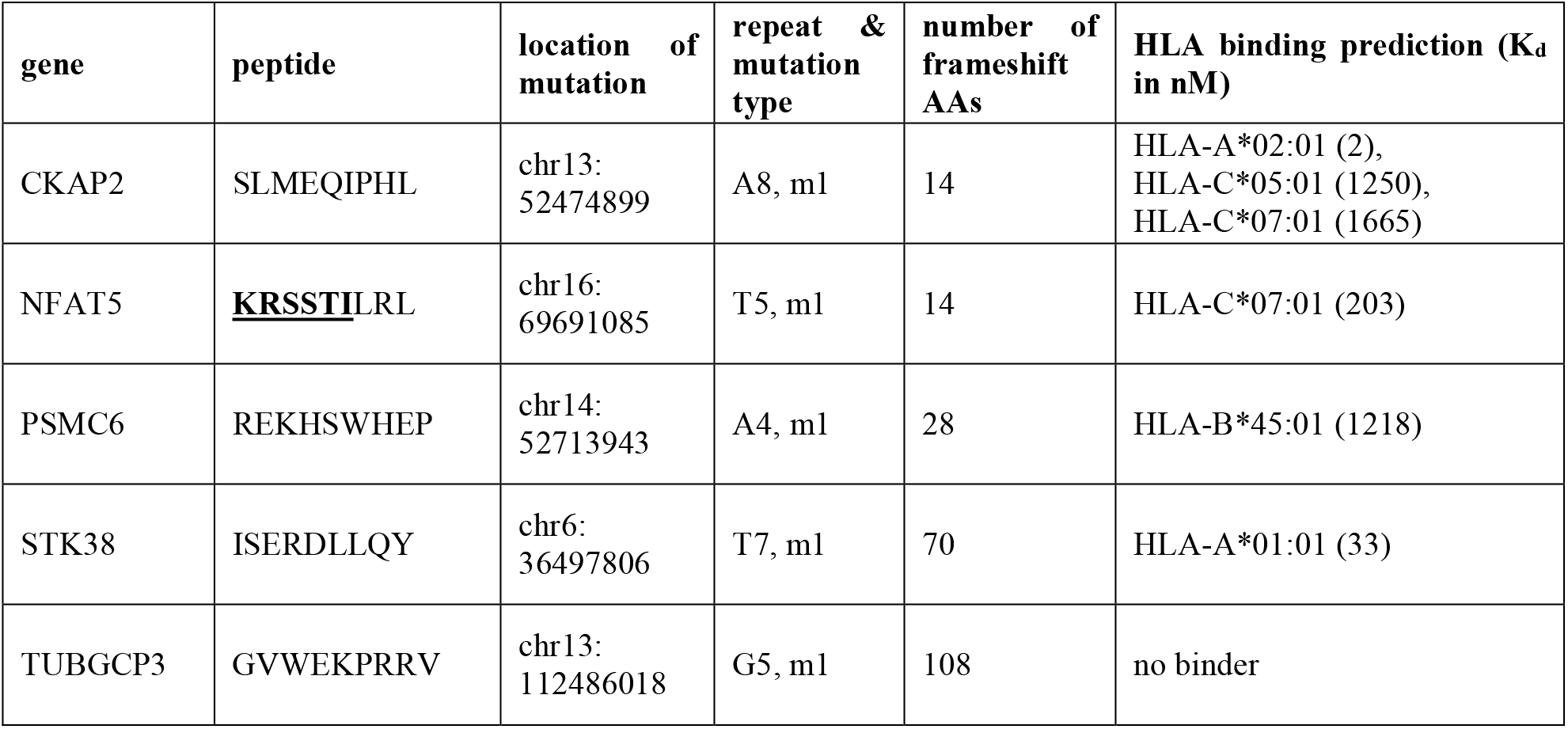
Overview of identified InDel neoepitopes. HLA binding prediction was performed with NetMHCpan 4.0. Underlined AAs of NFAT5 originate from wild-type NFAT5 protein sequence. m1, minus one base pair deletion.

| gene | peptide | location of mutation | repeat & mutation type | number of frameshift AAs | HLA binding prediction (K<sub>d</sub> in nM) |
| --- | --- | --- | --- | --- | --- |
| CKAP2 | SLMEQIPHL | chr13:<br>52474899 | A8, m1 | 14 | HLA-A*02:01 (2),<br>HLA-C*05:01 (1250),<br>HLA-C*07:01 (1665) |
| NFAT5 | <u>KRSSTILRL</u> | chr16:<br>69691085 | T5, m1 | 14 | HLA-C*07:01 (203) |
| PSMC6 | REKHSWHEP | chr14:<br>52713943 | A4, m1 | 28 | HLA-B*45:01 (1218) |
| STK38 | ISERDLLQY | chr6:<br>36497806 | T7, m1 | 70 | HLA-A*01:01 (33) |
| TUBGCP3 | GVWEKPRRV | chr13:<br>112486018 | G5, m1 | 108 | no binder |

**Table 2:**
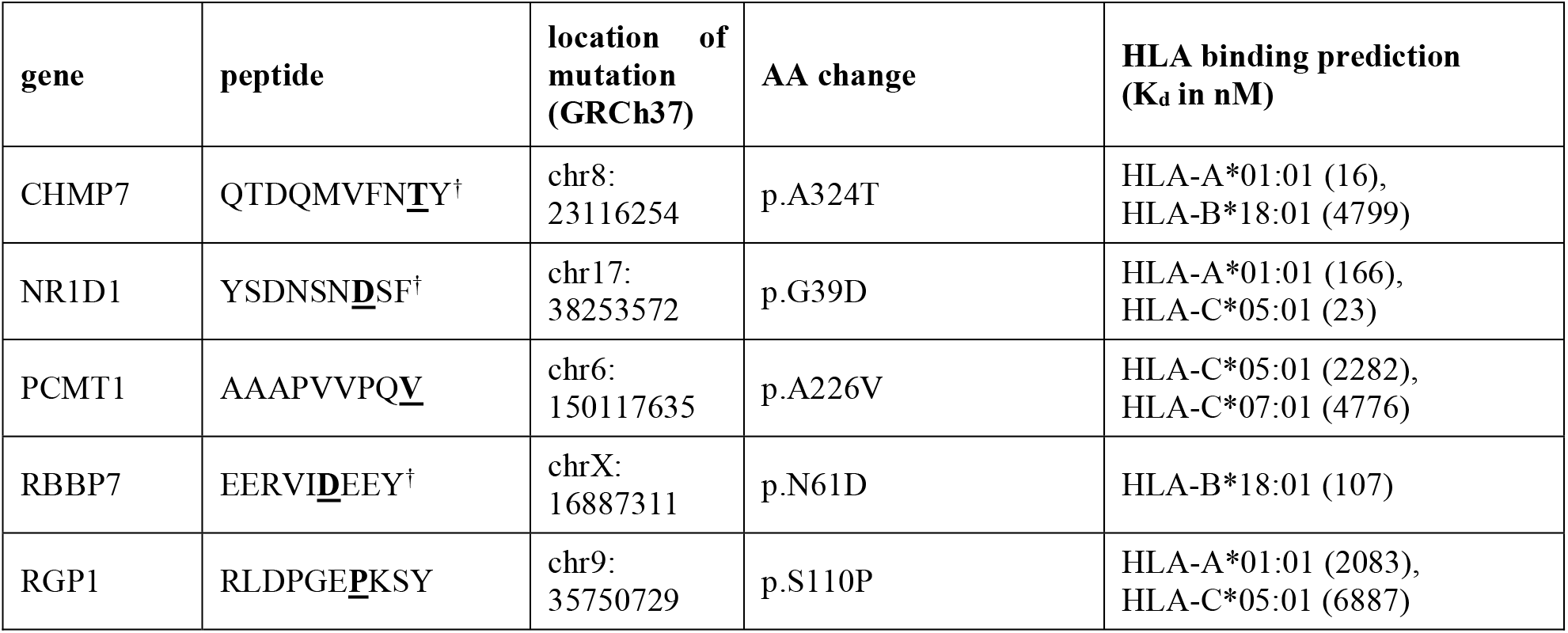
Overview of identified SNP neoepitopes. HLA binding prediction was performed with NetMHCpan 4.0. Mutated AAs originating from SNPs are underlined. Previously reported peptides (*26*) are marked with a dagger.

### CKAP2 frameshift mutation is recurrent in MSI CRC cell lines and patients

We next focused on the recurrence of frameshift mutations in the repeats of the five validated genes *CKAP2*, *NFAT5, PSMC6, STK38*, and *TUBGCP3* by analyzing 24 microsatellite-unstable colorectal cancer cell lines (Supplementary table 1). In addition to HCT-116, the *CKAP2* frameshift mutation was found in four other MSI CRC cell lines (KM12 (minus one base pair deletion (m1)), VaCo6 (m1), HROC24 (plus one base pair insertion (p1)), and LS411 (m1)). The *TUBGC3* and *STK38* frameshift mutations were identified in HCT-116 cells and in LoVo (m1) and HROC24 cells (m1), respectively. The *NFAT5* and *PSMC6* frameshift mutations were only found in HCT-116 cells. We next asked if the most recurrent *CKAP2* mutation could also be identified in MSI CRC patient samples. To this end, we analyzed genomic tumor DNA obtained from 56 MSI CRC patients and found m1 mutations in the described A8 repeat of the *CKAP2* gene in 9 samples (16%). Finally, we evaluated the potential InDel neoepitopes arising from the confirmed frameshift mutations. Binding prediction for overlapping nonamers originating from frameshifted protein sequences revealed multiple potential neoepitopes with promising binding affinities to common HLA supertypes (Supplementary figure 6)(*31–33*). Taken together, frameshift mutations were observed recurrently in cell lines derived from different tumors. *CKAP2* frameshifts emerged to be most interesting because these were found to be recurrent in cell lines and also in primary patient samples tested. Furthermore, the frameshifted CKAP2 protein sequence harbors several potential neoepitopes with binding potential to eight out of twelve HLA supertypes tested.

### NMD inhibition stabilizes frameshifted transcripts and augments HLA class I-mediated presentation of InDel neoepitopes

Apart from introducing frameshifts in the open reading frame and thus generating mRNAs encoding neoepitopes, InDel mutations typically trigger mRNA degradation by NMD (*16*). Therefore, the expression and consequently the presentation of most InDel neoepitopes in MSI CRC must be expected to be limited by NMD, reducing the usefulness of such neoepitopes for immunotherapy. We reasoned that NMD inhibition may stimulate the biosynthesis and the presentation of the InDel neoepitopes potentially increasing the cell’s visibility to the immune system. We have recently identified the licensed drug 5AZA as a pharmacologic inhibitor of NMD (*21*) and now tested the effect of this small molecule on the transcript and the peptidomic level in HCT-116 cells. First, we confirmed the known effect of 5AZA on NMD efficiency by measuring the abundance of known endogenous NMD targets using qPCR. The NMD target mRNAs *ATF3, ATF4, SC35C, SC35D,* and *UPP1* showed the expected upregulation between 1.8-fold and 4-fold following treatment with 5AZA (Figure 4A/Supplementary figure 1). We then analyzed the abundance of the frameshifted (FS) transcripts leading to the identified InDel neoepitopes. We found that 5AZA treatment highly significantly (*p* ≤ 0.0001) increased the abundance of *CKAP2* (2.0-fold), *PSMC6* (1.7-fold), *STK38* (1.4-fold), and *TUBGCP3* (1.9-fold) transcripts (Figure 4A). To further validate FS-transcripts as *bona fide* NMD targets, we measured mRNA levels after siRNA-mediated knockdown of the NMD core factor UPF1 (*13*). We observed significant increases in mRNA abundance in four out of the five targets (*CKAP2,* 1.4-fold; *PSMC6*, 1.5-fold; *NFAT5*, 1.4-fold; *TUBGCP3*, 1.6-fold; *p* ≤ 0.01)(Figure 4A).

**Figure 4:**
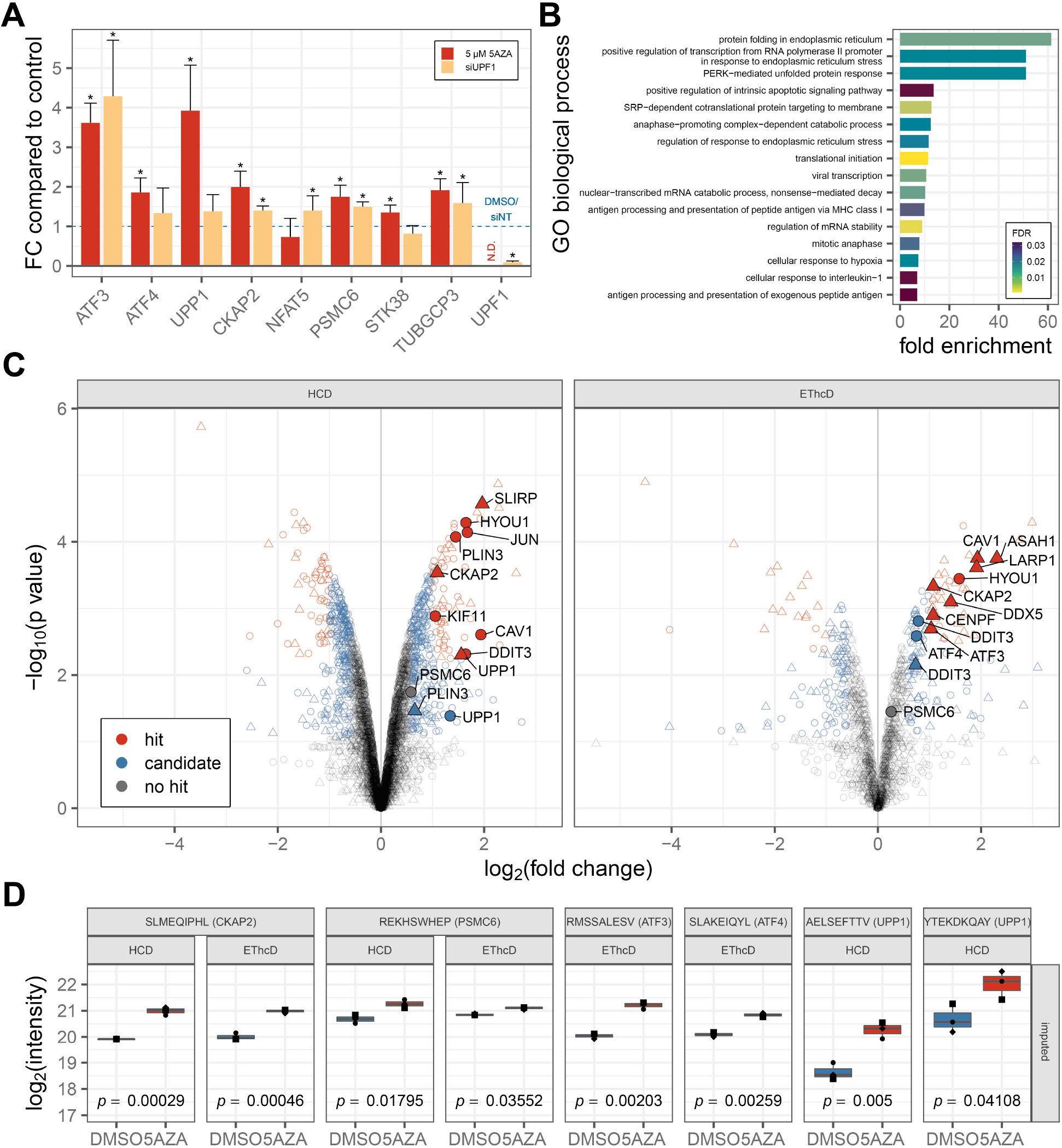
Treatment with 5AZA stabilizes NMD-targeted transcripts and augments HLA-mediated presentation of peptides originating from the encoded proteins. **(A)** qPCR analysis of endogenous NMD targets (*ATF3*, *ATF4*, *UPP1*) and InDel-mutated transcripts (*CKAP2, NFAT5*, *PSMC6, STK38*, *TUBGCP3*) after treatment with 5 µM 5AZA for 24 h (red) or siRNA-mediated KD of *UPF1* (orange). *UPF1* mRNA levels were determined as control for siRNA-mediated knockdown (N.D. = not determined). Each bar represents the mean ± SD of 3 experiments, \**p* ≤ 0.0001 (two-sided, unpaired t-test). **(B)** GO term enrichment for source genes of significantly upregulated hit peptides after 5AZA treatment for 24 h. **(C)** Volcano plot summarizing limma analysis of label-free quantification of the immunopeptidome isolated from 5AZA treated versus DMSO treated HCT-116 cells. Upregulated InDel neoepitopes (*CKAP2, PSMC6*) and peptides originating from putative endogenous NMD targets are labeled with the corresponding gene name. Color represents hit annotation, shape indicates if values were imputed (circle = no, triangle = yes). **(D)** Representative plots showing changes in intensity for InDel neoepitopes SLMEQIPHL (*CKAP2*), REKHSWHEP (*PSMC6*), and selected peptides originating from known NMD targets *ATF3*, *ATF4,* and *UPP1* after treatment with 5AZA for 24 h. Bars represent 25th to 75th percentiles, middle line represents median, points represent individual measurements of biological replicates.

We next tested if the increase of transcript levels by NMD inhibition is translated into an increased presentation of InDel neoepitopes at the cell surface and performed label-free quantification of HLA-presented peptides isolated from HCT-116 cells treated with either 5AZA or DMSO. Given the better correlation of raw intensity values between samples measured with the same method as well as the difference in the number of quantifiable peptides, the quantification workflow was performed separately for each MS fragmentation methodology (Supplementary figure 7A/B). We included only peptides that were measured in at least two out of three biological replicates per condition in the quantification dataset to minimize the need for imputing values. Data obtained with lowEThcD fragmentation were solely used for identification purposes and not for quantification. For the validated InDel neoepitopes, we complemented these untargeted data by integrating intensity values measured in a targeted MS2 analysis. Peptides were measured from the same samples and showed excellent reproducibility of intensities between untargeted and targeted MS2 data (Pearson’s correlation coefficient: 0.93; Supplementary figure 7C). After filtering, normalization, and imputation of missing values, the final quantification dataset consisted of 5072 distinct, quantifiable peptides (HCD: 4634 peptides, EThcD: 1376 peptides; overview of data processing in Supplementary figure 7D). Using the *limma* package, we identified a total of 838 differentially presented peptides upon NMD inhibition. 434 peptides showed an increased presentation while 404 were less abundant (Figure 4C). The composition of the immunopeptidome is known to be influenced by the composition of the proteome (*26, 34*). We thus validated the effect of NMD inhibition on the immunopeptidome by confirming that the GO term categories of the source proteins of differentially presented HLAp mirror those known to be affected by NMD inhibition at the proteome level (*35*). GO term enrichment analysis for source genes of upregulated hit peptides following NMD inhibition revealed these to be involved in protein folding, ER stress response, unfolded protein response, proteasome-mediated APC-dependent catabolic process (*i.e.* breakdown of proteins by peptide bond hydrolysis) as well as antigen processing and presentation (Figure 4B). The effect of 5AZA treatment on the presentation of peptides originating from known NMD targets was confirmed by the significant upregulation of several *ATF3*, *ATF4,* and *UPP1* peptides (Figure 4D). We further validated the presentation of peptides originating from endogenous NMD targets by comparing source proteins of identified peptides with previously reported and ENSEMBL annotated NMD targets (*16, 36–38*). This analysis revealed several peptides originating from *ASAH1*, *CAV1*, *DDIT3*, *DDX5*, *HYOU1*, *JUN*, *PLIN3*, and *SLIRP* to be upregulated in the immunopeptidome of HCT-116 cells following NMD inhibition by 5AZA. We next studied endogenous peptides originating from the non-frameshifted 5’ sequences of FS-bearing transcripts. This analysis revealed peptides originating from *CENPF*, *KIF11*, and *LARP1* to be upregulated following NMD inhibition by 5AZA (Figure 4C). Finally, we analyzed the effect of 5AZA treatment on the presentation of the validated InDel neoepitopes. The HLA class I-mediated presentation of the *CKAP2*-derived InDel neoepitope was significantly upregulated 2.1 – 2.13-fold (*p* ≤ 0.0005) and that of the *PSMC6*-derived InDel neoepitope significantly (*p* ≤ 0.05) albeit less strongly (1.2 – 1.5-fold) upregulated both in the HCD and the EThcD datasets (Figure 4D). Taken together, these findings show that modulation of NMD efficiency in MSI CRC cells by the pharmacologic NMD inhibitor 5AZA stabilizes FS-bearing and NMD-targeted transcripts and results in the increased cell surface presentation of HLAp derived thereof.

### *In vivo* immunization with InDel neoepitopes induces CD8^+^ T cell responses

We next analyzed the potential of InDel neoepitopes to induce specific T cell responses by performing *in vivo* immunizations in a humanized HLA-A*02:01-transgenic mouse model (*39*). Binding predictions indicated that the *CKAP2*-derived InDel neoepitope is a strong HLA-A*02:01 binder (percentile rank: 0.0071, predicted IC50: 2.3736 nM) while InDel neoepitopes derived from *NFAT5, PSMC6*, and *STK38* were predicted to bind other alleles than HLA-A*02:01 (Table 1). Interestingly, the *TUBGCP3*-derived InDel neoepitope was not predicted to bind any of the HLA alleles expressed by HCT-116 cells by NetMHCpan but showed the strongest affinity to HLA-A*02:01 (percentile rank: 2.3267, predicted IC50: 6217 nM). We have therefore included this InDel neoepitope for further testing.

As a first step, we immunized three mice with a mixture of peptides consisting of two potential HLA-A*02:01 binders (*CKAP2*- and *TUBGCP3*-derived InDel neoepitopes) and two “non-binders” (*NFAT5*- and *PSMC6*-derived InDel neoepitopes) as negative controls. Whole splenocytes were analyzed by *ex vivo* IFNγ ELISpot assays. One out of three mice generated a peptide-specific T cell response against the *CKAP2*-derived InDel neoepitope and two out of three mice generated a peptide-specific T cell response against the *TUBGCP3*-derived InDel neoepitope. As expected, immunization did not induce peptide-specific T cells for predicted “non-binder” InDel neoepitopes derived from *NFAT5* and *PSMC6* (Supplementary figure 8). These results were validated by immunizations of mice with a single peptide according to the immunization scheme shown in Figure 5A. The analysis of isolated splenic CD8^+^ T-cells following immunizations with either *CKAP2*- or *TUBGCP3*-derived InDel neoepitopes or the positive control HPV16 peptide E7 (AAs 11-19) resulted in IFNγ-specific and highly significant responses in the ELISpot assay (Figure 5B/C). These data demonstrate that InDel neoepitopes are processed and presented *in vivo* via HLA-A*02:01 molecules. Importantly, immunization can induce a specific CD8^+^ T cell-mediated immune response.

**Figure 5:**
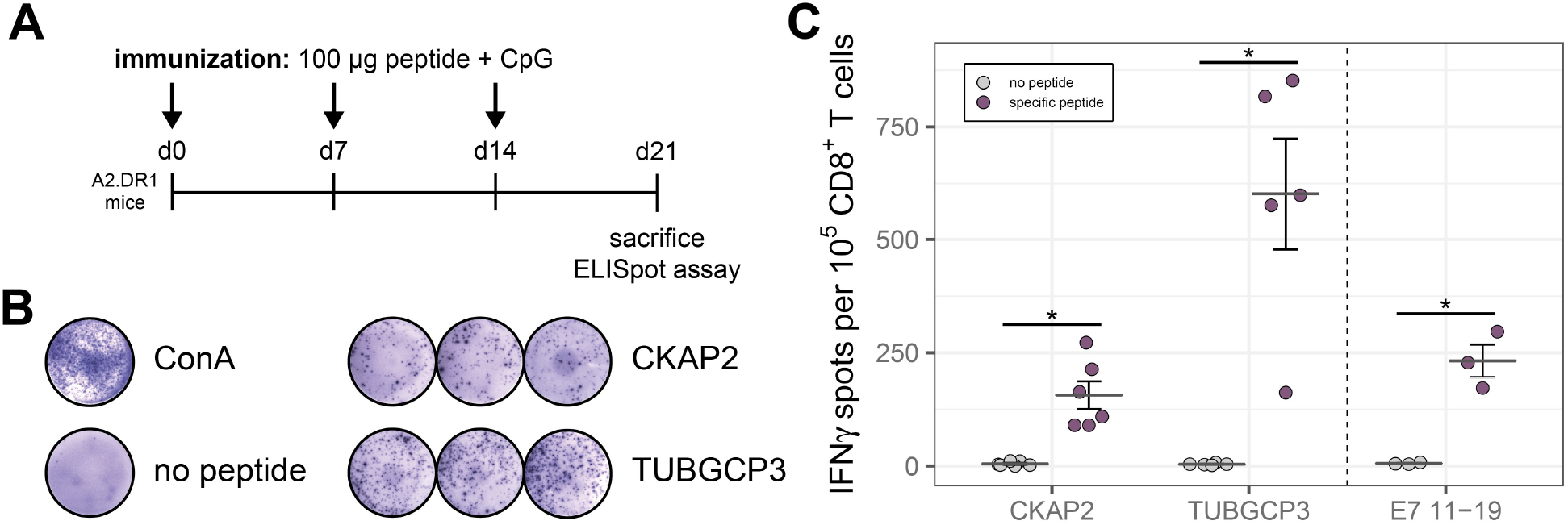
In vivo immunization of A2.DR1 mice with InDel neoepitopes induces CD8^+^ T cell responses. **(A)** Immunization scheme. **(B)** Representative ELISpot assay results for isolated CD8^+^ T cells stimulated with ConA (assay positive control), no peptide control, and InDel neoepitopes SLMEQIPHL (*CKAP2*) and GVWEKPRRV (*TUBGCP3*). **(C)** Quantitative analysis of ELISpot assays. Bars represent mean ± SEM of N = 6 (*CKAP2*), N = 5 (*TUBGCP3*) or N = 3 (E7 11-19, control peptide) experiments, \**p* ≤ 0.005 (two-sided, unpaired t-test).

## Discussion

Identification of tumor-specific neoepitopes represents a crucial step in the development of therapeutic cancer vaccines and a high load of neoepitopes has been associated with effective immunotherapy (*5*). While *in silico* predictions have been employed previously to identify cancer-specific neoepitopes, only a negligible fraction of candidates implicated by this approach are actually presented by HLA class I molecules. Therefore, most candidates are unable to elicit anti-tumor immune responses (*40*). Mass spectrometry-based interrogation of the immunopeptidome provides an unbiased view of actually presented peptides. Previous studies of neoepitopes using mass spectrometry mainly focused on SNP-derived neoepitopes (*26*), however, SNP-derived neoepitopes are thought to induce less robust immune responses than InDel-derived neoepitopes (*6*). Other studies have analyzed only InDel neoepitopes originating from specific, recurrent InDel mutations (*41, 42*), or failed to detect FS-derived mutant sequences at the proteome level (*43*). Here, we report the first unbiased, mass spectrometry-based identification of immunogenic InDel neoepitopes using publicly available sequencing data. We demonstrate that inhibition of nonsense-mediated mRNA decay stabilizes the corresponding FS-bearing mRNAs consequently increasing HLA class I-mediated presentation of InDel neoepitopes at the cell surface.

We combined a previously established high-throughput procedure for immunoprecipitation of HLA:peptide complexes (*27*) with different fragmentation and precursor selection methods for LC-MS/MS to obtain a representative view of the immunopeptidome of the MSI CRC cell line HCT-116. While samples measured using HCD fragmentation yielded the majority of HLAp identifications due to the higher MS2/MS1 rate and therefore deeper sampling of the immunopeptidome compared to dual-fragmentation measurements, lowEThcD fragmentation identified a subset of low-abundance HLAp, which were found neither by EThcD with standard precursor selection nor by HCD. Therefore, lowEThcD (or any other methodology targeting low-abundance precursors first) is suited to further expand the detectable HLA class I immunopeptidome.

Next, we performed a multi-round database search, matching high-scoring “*de novo* only” spectra, which did not match any known human protein sequences, against custom InDel and SNP neoepitope databases and identified 14 neoepitope candidates. While eleven of these candidates were identified using standard HCD fragmentation, three could only be identified using EThcD or lowEThcD fragmentation, emphasizing the added value of combining different fragmentation methods. As an important technical conclusion, our work thus demonstrates that the combination of different fragmentation and precursor selection methodologies can increase the number of identified HLA class I-presented (neo)epitopes.

It is important to note that the number of neoepitopes identified by our approach may be limited. Variant calling of InDel mutations remains challenging to date (*44*) and although the usage of public mutation databases is accepted as a reasonable substitute for sequencing (*18*), it must be considered that the data used for the construction of our InDel neoepitope databases might be incomplete or biased. Indeed, the length of mutated microsatellites resulting in identified InDel neoepitopes was rather short (4 – 8 bp) and we did not document InDel neoepitopes derived from well-established InDel mutations occurring in longer repetitive sequences of the HCT-116 cell line as the mutations are not documented in either the COSMIC or CCLE database. To address this limitation we also searched our MS raw data against a database of frequent InDel mutations in MSI CRC (*44*), although this did not yield new InDel neoepitope identifications. In future studies, this limitation may be overcome and more InDel neoepitopes might be identified by utilizing long-read sequencing technologies (*45*). Moreover, we did not identify the known HLA-A*02:01-restricted *TGFBR2*-derived InDel neoepitope RLSSCVPVA (*7, 46*) in our targeted MS approach. Based on a recently reported analysis of T cell responses against this InDel neoepitope, this negative result is likely explained by the low stability of the HLA class I:peptide complex (*47*) which leads to the peptide being overlooked in approaches involving immunopurification of HLA molecules.

As a second important finding, we report here that NMD inhibition with 5AZA significantly augments HLA class I-mediated presentation of peptides originating from NMD-sensitive transcripts, including InDel neoepitopes, thus conceptually increasing the likelihood of the recognition of tumor cells by the immune system. We found the m1 mutation resulting in the *CKAP2*-derived neoepitope to be highly recurrent in MSI CRC cell lines and patient samples thus confirming the frequent mutation of this gene in MSI CRC patients (*48*). While the FS-bearing *CKAP2* transcript has previously been classified as likely being NMD-resistant based on the localization of the m1 mutation (*49*), we directly demonstrate NMD sensitivity of this transcript by pharmacological NMD inhibition with 5AZA and by RNAi of the key NMD factor UPF1. In agreement with previous reports (*50, 51*) these findings indicate that sequence features alone are not sufficient to predict NMD sensitivity and require experimental validation. Notably, in addition to the FS-bearing NMD targets, many HLA-presented peptides whose HLA class I-mediated presentation was upregulated following 5AZA originated from source proteins involved in stress response mechanisms. These findings confirm our previously reported data showing that many stress-related transcripts are controlled by NMD, and support the hypothesis that NMD inhibition augments the expression of physiologic C-terminally truncated proteins (*35*).

Finally, we show directly in a humanized HLA-A*02:01-transgenic mouse model that the identified HLA-A*02:01-restricted InDel neoepitopes can induce strong CD8^+^ T cell responses, demonstrating their *in vivo* processing, presentation, and immunogenic potential. InDel neoepitopes identified by this approach thus might serve as valuable starting points for the development of a vaccine or engineered T cell therapies. As InDel neoepitopes are encoded by frameshifted transcripts that in many cases are NMD targets, pharmacologic inhibition of NMD by the clinically approved drug 5AZA may act synergistically with immune checkpoint inhibition. Moreover, since frameshifted transcripts often encode multiple InDel neoepitopes, we envision that NMD inhibition could increase HLA class I-mediated presentation of InDel neoepitopes independently of a patient’s HLA genotype. While we have chosen MSI colorectal cancer as the proof-of-concept model disease, microsatellite instability is increasingly recognized in several other malignancies which might benefit from increased neoepitope presentation after NMD inhibition (*52*). In summary, we here show the feasibility of high-throughput immunopeptidomics for the identification of immunogenic InDel neoepitopes in the context of MSI and provide evidence that pharmacological NMD inhibition augments their HLA class I-mediated presentation, thus turning cancer cells into easily identifiable targets for tumor-specific T cells.

## Materials and Methods

### Mouse strain

The HLA-A2.1/HLA-DR1-transgenic H-2 class I-/class II-knockout mice (*39*) were provided by the Institute Pasteur (Paris, France). All animal procedures followed the institutional laboratory animal research guidelines and were approved by the governmental authorities. The mice were fed a standard chow diet and provided water *ad libitum*. The Animal Care Facilities at DKFZ have been approved by FELASA and accredited. For the experiments, mice were assigned to age-matched and sex-matched groups.

### Human Tissues

Human tissues were obtained from the local tissue bank established within the German Collaborative Group on HNPCC. Informed consent was obtained from all patients and the study protocol was approved by the local Ethics Committee (S-583/2016). For all tissue samples, MSI status has been determined previously based on the National Cancer Institute/ICGHNPCC reference marker panel (*53*) and CAT25 as an additional mononucleotide marker (*54*). MSI is defined by instability in at least 30% of tested markers.

### Cell lines

All human CRC cell lines have been described previously (*55–57*). HCT-116 cells (ATCC® CCL-247™) cells were maintained in RPMI1640 medium supplemented with 10% FCS and 1% P/S. HLA-I types were previously determined as HLA-A*01:01, HLA-A*02:01, HLA-B*18:01, HLA-B*45:01, HLA-C*05:01, and HLA-C*07:01 (*22*) and confirmed by sequencing in the DKMS Life Science Lab GmbH (Dresden, Germany). HB95 cells (ATCC® HB-95™) were maintained in CELLine bioreactor flasks in RPMI1640 medium supplemented with 10% FCS and 1% P/S. NMD efficiency of HCT-116 cells was assessed in triplicates using a previously published NMD reporter system (*23*). All cell lines were tested negative for mycoplasma contamination.

### High-throughput purification of HLA class I-peptides

Antibody purification and coupling were performed as described previously (*58*). *In vitro* treatment was performed in biological triplicates. Briefly, 7.3×10^6^ HCT-116 cells were seeded per 150 mm dish and after 48 h, cells were treated either with a final concentration of 5 µM 5AZA (Sigma-Aldrich) or with DMSO (negative control) for 24 h. Cells were harvested by scraping in cold PBS and aliquots of 1×10^8^ cells were stored as snap-frozen dry pellets at −20°C. Snap-frozen dry pellets were lysed immediately before IP. IP of HLA class I:peptide complexes and separation of HLAp was performed as previously described (*27*) omitting pre-clear and HLA class II plates.

### LC-MS/MS analysis

For LC-MS/MS analysis pooled samples were resuspended in 30 µl of 0.1% formic acid and 4.5 µl were used per injection. Lyophilized synthetic peptides for validation of neoepitopes were purchased from JPT Peptide Technologies (Berlin, Germany) and diluted to a concentration of 100 fmol/ul and 3 µl were used per injection. The mass spectrometric analysis was conducted using an UltiMate™ 3000 RSLCnano system (Thermo Fisher Scientific) directly coupled to an Orbitrap Fusion Lumos (Thermo Fisher Scientific). Peptides were loaded onto the trapping cartridge (µ-Precolumn C18 PepMap 100, 5µm, 300 µm i.d. x 5 mm, 100 Å) for 3 min at 30 µL/min (0.05% TFA in water). Peptides were eluted and separated on an analytical column (nanoEase MZ HSS T3 column, 100 Å, 1.8 μm, 75 μm x 250 mm) with a constant flow of 0.3 µL/min using solvent A (0.1% formic acid in LC-MS grade water) and solvent B (0.1% formic acid in LC-MS grade acetonitrile). Total analysis time for the HCD method was 90 min with a gradient containing an 8-25% solvent B elution step for 69 min, followed by an increase to 40% solvent B for 5 min, 85% B for 4 min and re-equilibration step to initial conditions. The LC system was coupled online to the mass spectrometer using a Nanospray-Flex ion source (Thermo Fisher Scientific) and a Pico-Tip Emitter 360 µm OD x 20 µm ID; 10 µm tip (New Objective). The MS was operated in positive mode and a spray voltage of 2.4 kV was applied for ionization with an ion transfer tube temperature of 275 °C. Full scan MS spectra were acquired in profile mode for a mass range of 300-1650 *m/z* at a resolution of 120 000 (RF Lens 30%, AGC target 4e5 ions, and maximum injection time of 250 ms). The instrument was operated in data-dependent mode for MS/MS acquisition. Peptide fragment spectra were acquired for charge states 1-4. The quadrupole isolation window was set to 1.2 m/z and peptides were fragmented via HCD (30%). Fragment mass spectra were recorded at a resolution of 30 000 for a maximum of 2e5 ions (AGC target) or after 150 ms maximum injection time. The instrument acquired MS/MS spectra for up to 3 s between MS scans. Dynamic exclusion was set to 20 s. Additionally, samples were analyzed using two HCD/EThcD decision tree methods. Here, the instrument fragmented precursors with a charge state of +1 using the parameters of the HCD method. Charge states 2-7 were fragmented using ETD (Calibrated Charge-Dependent ETD Parameters) with supplemental activation enabled (HCD, 30%). For MS/MS spectra acquisition, either high abundant (EThcD) or low abundant (lowEThcD) precursors were selected first. AGC target was set to 2e5 ions and a maximum injection time of 200 ms was allowed and the resulting MS/MS spectra were recorded in the Orbitrap with a resolution of 30 000. Total analysis time for the HCD/EThcD decision tree methods was 180 min with a gradient containing an 8-25% solvent B elution step for 150 min, followed by an increase to 40% solvent B for 14 min, 85% B for 4 min and re-equilibration step to initial conditions. To further validate the presence of the candidates and to obtain reliable quantification data, InDel neoepitope candidates were measured from the same samples using a targeted MS2 method with the previously described settings. Precursor masses of targeted InDel neoepitope candidates are listed in Supplementary table 2 and were fragmented using both HCD and EThcD.

### Generation of neoepitope databases

Frameshift peptide databases were constructed based on publicly available sequencing information from CCLE and COSMIC for HCT-116 using in-house developed R scripts. Published frameshift sequences originating from recurrent InDel mutations in MSI CRC were included in a separate database (*44*). Previously published information on potential neoepitopes resulting from single nucleotide polymorphisms were included in a separate database (*22*).

### MS data analysis and identification of HLAp

Mass spectrometry raw data were analyzed using PEAKS Studio X (version 10.0, Bioinformatics Solutions Inc.). Raw files were subjected to the default data refinement before *de novo* sequencing and database search. The parent mass error tolerance was set to 10.0 ppm while the fragment mass error tolerance was set to 0.02 Da. Fragmentation mode was set either to “HCD” or “Mixed” (for EThcD measurements with HCD fragmentation for single positive precursors). All raw files were first searched against the UniProt/SwissProt database (20659 entries, February 2019) and a database containing standard contaminants with oxidation of methionine (15.99 Da), carbamidomethylation of cysteine (57.02 Da), and acetylation of N-termini (42.01) as variable modifications. The enzyme specificity was set to “no enzyme”. *De novo* only spectra with an average local confidence of more than 50% (*i.e.* good spectra not matching a UniProt/SwissProt database entry), were subjected to a multi-round database search using the in-house generated frameshift peptide databases based on CCLE (747 entries), COSMIC (1071 entries) and the SNP neoepitope datasets (1526 entries). All peptides identified at a peptide spectrum match FDR of 1% were exported and contaminants, as well as peptides shorter than 8 AAs and longer than 15 AAs, were filtered out. Unique peptides matching to entries of the UniProt/SwissProt database were reported as “endogenous”, while peptides matching one of the frameshift peptide databases were reported as “InDel neoepitopes” and peptides matching the SNP neoepitope database were reported as “SNP neoepitopes”. HLAp were verified *in silico* and label-free quantification was performed using an in-house developed R script (see Supplementary Materials and Methods for details).

### Mutation analysis

Underlying frameshift mutations of InDel neoepitopes and single nucleotide polymorphisms of SNP neoepitopes were verified by Sanger sequencing of PCR amplified fragments (Eurofins Genomics Germany GmbH, Köln) and analyzed using the Indigo webtool (*59*). Mutation analysis of MSI CRC cell lines and patient samples was performed using fluorescently labeled primers for amplification. Fragments were visualized on an ABI3130xl (Applied Biosystems) genetic analyzer as described previously (*54*). Primers are reported in Supplementary tables 3-5.

### Quantitative real-time PCR

RNA was isolated using TriReagent (Sigma). 2 µg of RNA were reverse transcribed using the Revert-Aid™ H Minus Reverse Transcriptase Kit (Thermo Scientific) according to the manufacturer’s protocol. qPCR was performed in technical triplicates on a StepOnePlus™ system (Applied Biosystems) using primaQuant CYBR qPCR Master Mix (Steinbrenner Laborsysteme). Primers for *ATF3* (NMD-sensitive), *ATF4* (NMD-sensitive), *SC35A* (NMD-insensitive control), *SC35C* (NMD-sensitive), *SC35D* (NMD-sensitive), *UPP1* (NMD-sensitive), and the house-keeping gene *HPRT1* were reported elsewhere (*21, 60, 61*). qPCR primers for neoepitope candidates are reported in Supplementary table 6.

### Immunization and ex vivo IFNγ ELISpot assay

HLA-A2.1/HLA-DR1-transgenic H-2 class I-/class II-knockout mice (*39*) were immunized weekly for three weeks with either a peptide pool consisting of 50 µg of *CKAP2*, *NFAT5*, *PSMC6*, and *TUBGCP3* peptide each or 100 µg of *CKAP2* or *TUBGCP3* peptides separately purchased from JPT Peptide Technologies (Berlin, Germany) or synthesized by the GMP & T Cell Therapy Unit at German Cancer Research Center (DKFZ; Heidelberg, Germany) and 20 µg CpG ODN 1826 (TIB MolBiol) suspended in 20 µl PBS. IFNγ ELISpot was performed *ex vivo* seven days after the last immunization with isolated splenocytes or CD8^+^ T cells as described previously (*44*). CD8^+^ T cells were isolated with the CD8a^+^ T Cell Isolation Kit (Miltenyi Biotec) following the manufacturer’s instruction using LS columns.

### Statistical analysis

Luciferase assay and qPCR data are presented as mean *±* SD from three independent experiments. ELISpot assay data are presented as mean *±* SEM scatter dot plots from three to six independent experiments. Statistical analyses were made using a two-sided, unpaired t-test with correction for multiple hypothesis testing. *p* ≤ 0.05 was considered significant.

## Acknowledgments

We thank Michal Bassani-Sternberg (University of Lausanne, Switzerland) for providing the HB-95 hybridoma cell line. We thank Vera Fuchs and Jonathan Dörre for their excellent technical assistance. We are grateful to Beate Amthor, Michael Backlund, Annika Brosig, Manouk Gerritsen, Claudia Gruber, Margit Happich, Daria Lavysh, and Gabriele Tolle for their support. *Funding*: J.P.B. receives a Ph.D. scholarship by the Fastenrath-Stiftung. A.E.K. is supported by the Manfred-Lautenschläger-Stiftung. *Author contributions*: J.P.B., M.W.H., and A.E.K. designed the project. J.P.B. performed most of the experiments, performed the data analysis, generated the figures, and drafted the manuscript. D.H. contributed to the experimental design and establishment of MS methodologies. D.H. and M.R. performed MS runs. F.S. contributed to the quantitative MS data analysis. K.U. and A.H.-S. performed in vivo immunizations and ELISpot assays. J.G., M.K., G.N.-Y., and M.v.K.D. contributed to the conceptualization of the study and interpretation of the results. M.W.H. and A.E.K. coordinated the study and finalized the manuscript with input from all authors. *Competing interests*: All authors declare that they have no competing interests. *Data and materials availability*: The MS data have been deposited to the ProteomeXchange Consortium via the PRIDE partner repository (*62*) with the dataset identifier PXD021755. Scripts are available upon request.

## Supplementary Materials

### Supplementary Materials and Methods

#### In silico methods for validation of HLAp

Binding prediction to the expressed HLA allotypes was performed using NetMHCpan (version 4.0a)(*63*). The rank threshold for binders was set to <0.5% for strong binders (SB) and <2% for weak binders (WB). For peptides binding to more than one HLA allotype, only the best ranked HLA allotype with its corresponding affinity in nM was reported. Peptide sequences were clustered using GibbsCluster (version 2.0)(*64*) with default parameters for MHC class I ligands and motifs were visualized with Seq2Logo (version 2.0)(*65*). Hydrophobicity indices of identified peptides were calculated using SSRcalc version Q (*28*) using the following parameters: 100Å C18 column, 0.1% Formic Acid (2015 model), no cysteine protection. Spectra recorded from synthetic peptides were compared to spectra measured from the samples. Briefly, all spectra recorded for a given peptide were compared to all spectra recorded from its synthetic counterpart. Raw data for matched fragment ions was exported from PEAKS Studio X (PSM-ions.txt), normalized intensities of matched fragment ions were correlated and the match with the highest correlation was reported graphically. Moreover, retention time differences between the two peptides were calculated and reported.

#### Label-free quantification of HLAp

Quantification of HLA-presented peptides was performed using the raw output (peptides.csv) from PEAKS and a custom script in the R programming language (ISBN 3-900051-07-0). First, measured intensities from InDel neoepitope candidates using parallel reaction monitoring were combined with measured intensities from endogenous peptides, and data were filtered to include only peptides which were measured in at least two out of three replicates per condition and MS method used. Subsequent steps were performed for each MS method separately. Potential batch effects between biological replicates were removed using the *limma* package (*66*). Next, the variance stabilization normalization method was performed on the log2 transformed raw data using the vsn package (*67*) with individual normalization coefficients for the different MS methods. Missing values were imputed with the Msnbase package using nearest neighbor averaging (*68*). Normalized data were tested for differential HLA presentation of peptides between DMSO and 5AZA treated samples using the *limma* package. The replicate factor was included in the linear model. Peptides with an FDR ≤ 0.05 and an FC > 2 were defined as hits while peptides with an FDR ≤ 0.2 and an FC ≥ 1.5 were defined as candidates. GO-term analysis was performed for hits with the PANTHER classification system (*69*) using the following parameters: PANTHER overrepresentation test with “*Homo sapiens* (all genes in database)” as reference, “GO biological process complete” as annotation dataset, and Fisher’s Exact with FDR correction for multiple testing. Results were visualized using the ggplot2 package (ISBN 978-3-319-24277-4).

### Supplementary figures

**Supplementary figure 1:**
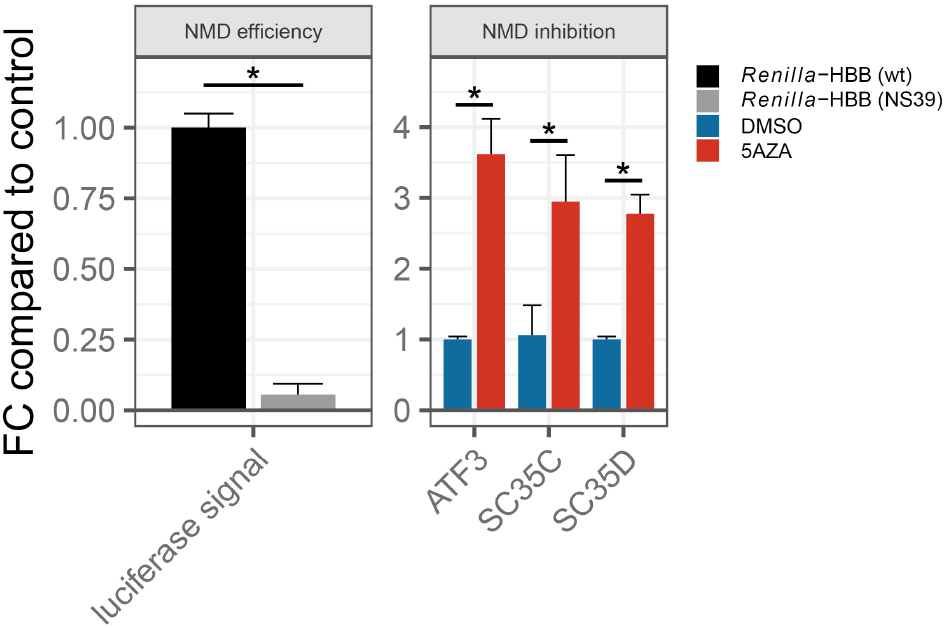
NMD is efficient in HCT-116 cells and can be inhibited by 5AZA. NMD efficiency determined by significant downregulation of luciferase signal from *Renilla*-HBB (NS39) construct compared to *Renilla*-HBB (wt) construct (left panel). Each bar represents mean ± SD of 3 experiments, \**p* ≤ 0.0001 (two-sided, unpaired t-test). Treatment with 5 µM 5AZA for 24 h increases mRNA abundance of endogenous NMD targets *ATF3*, *SC35C*, and *SC35C* as determined by qPCR (right panel). Each bar represents mean ± SD of 3 experiments, \**p* ≤ 0.0001 (two-sided, unpaired t-test).

**Supplementary figure 2:**
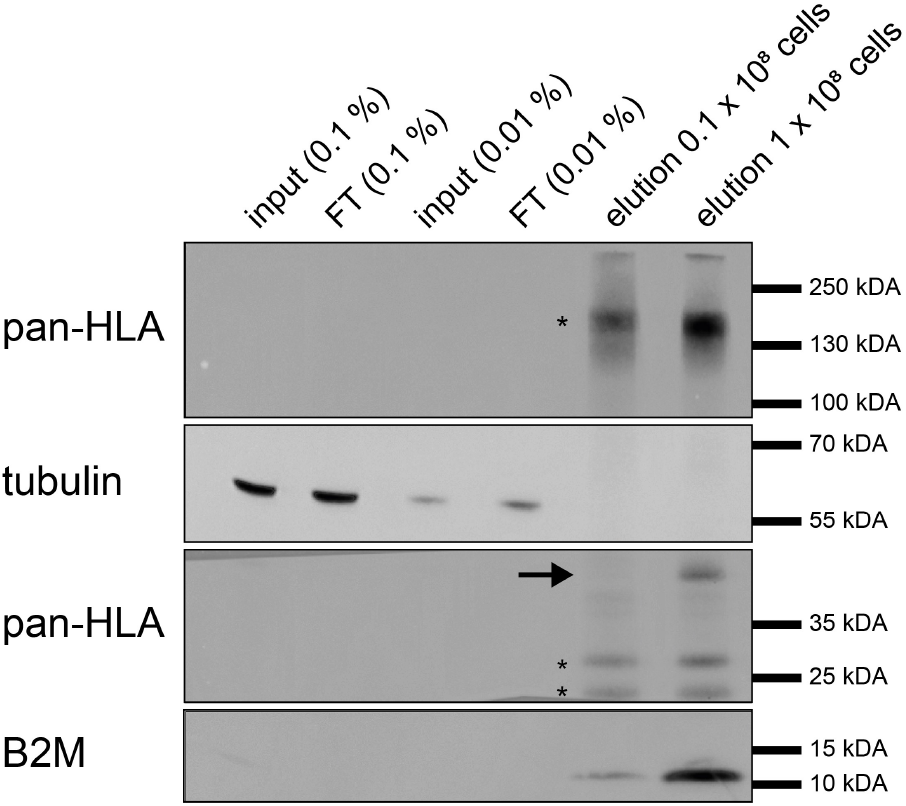
Selective purification of HLA molecules by high-throughput immunoprecipitation. Western blot analysis of HLA molecules purified by immunoprecipitation with W6/32 antibody. Arrow indicates expected band of HLA molecules. Asterisks indicate background bands of eluted W6/32 antibody. B2M, beta-2-microglobulin.

**Supplementary figure 3:**
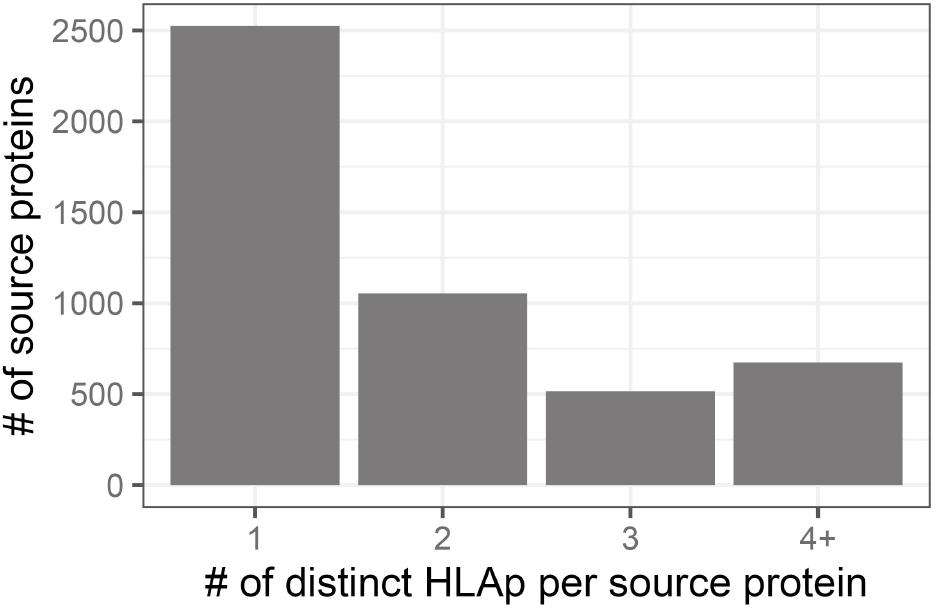
Most source proteins are represented only by one HLAp. Number of distinct identified HLAp per source protein.

**Supplementary figure 4:**
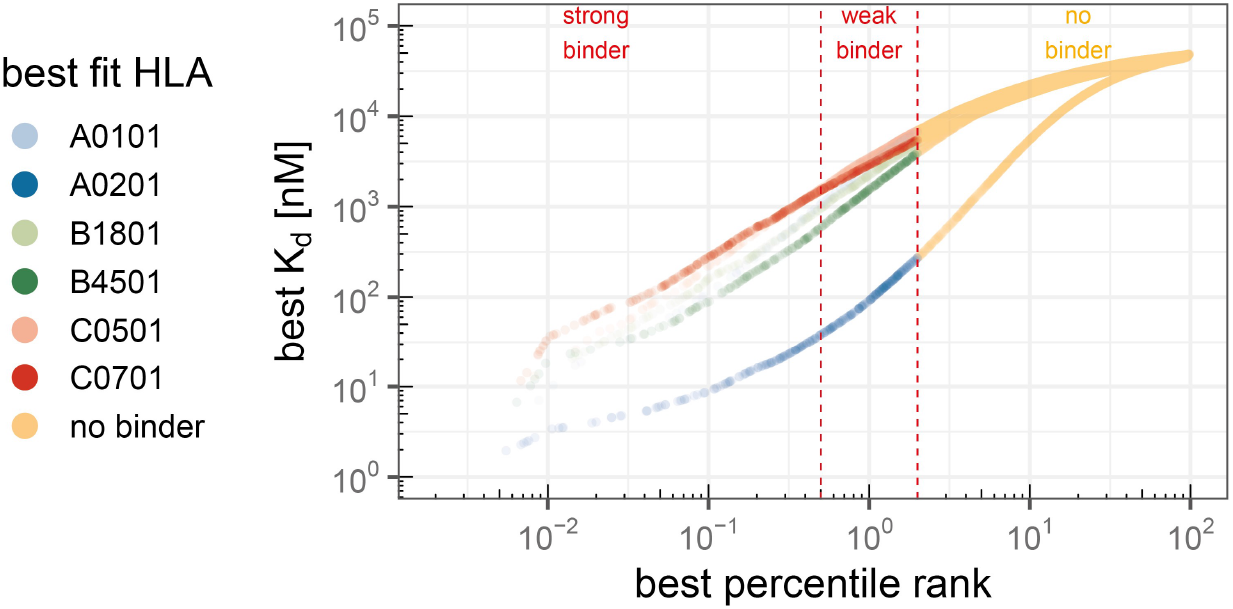
Frameshift inducing InDel mutations generate 2782 potential InDel neoepitopes. Binding prediction of all potential nonamers resulting from InDel mutations annotated in COSMIC and CCLE databases to HLA alleles expressed by HCT-116 cells. Threshold for strong binders is top 0.5% ranked, for weak binders top 2%.

**Supplementary figure 5:**
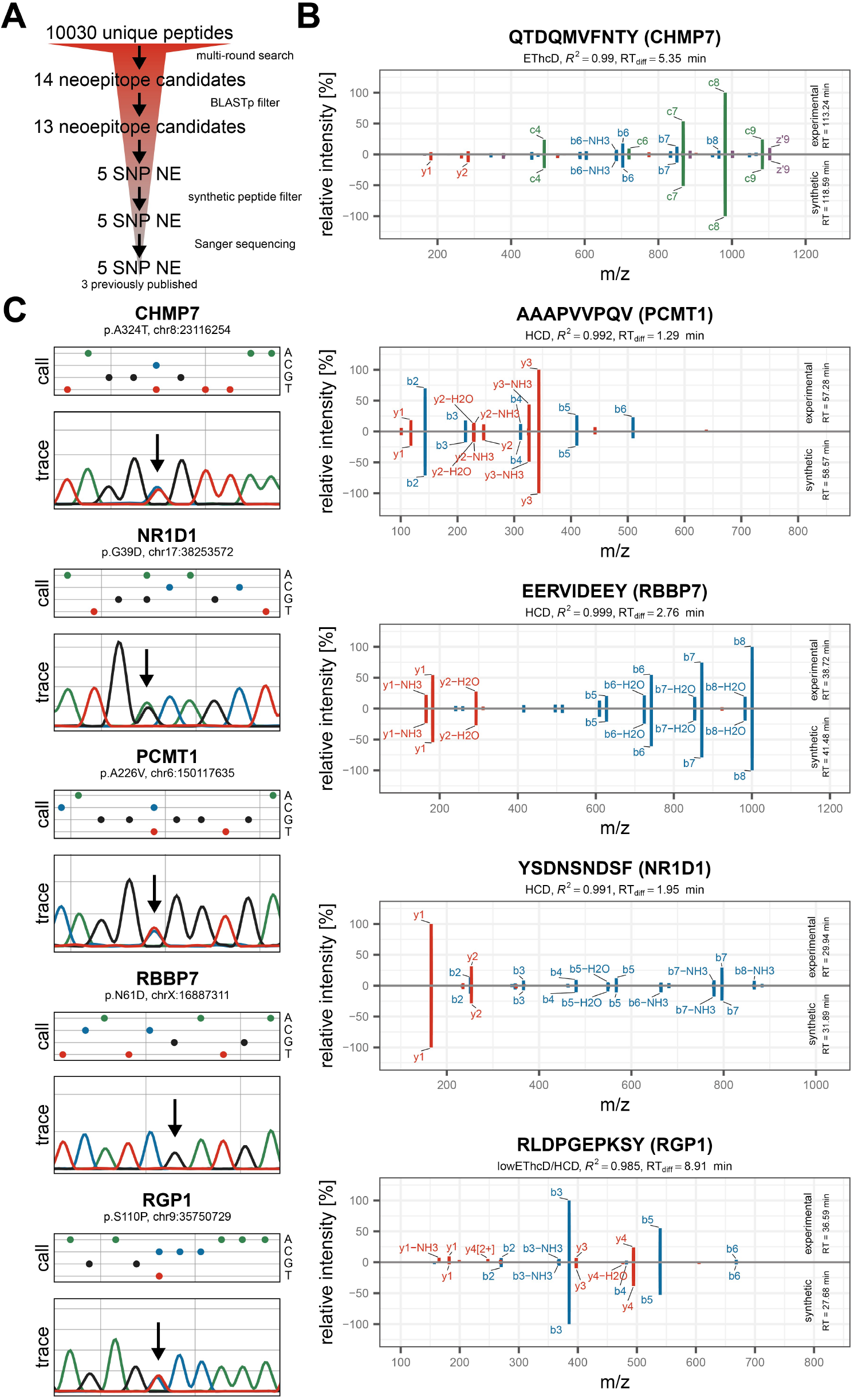
Validation of identified SNP neoepitopes. **(A)** Overview of validation procedure. Candidates were filtered using BLASTp to exclude peptides matching endogenous proteins. Spectra of candidates were compared to spectra recorded from synthetic peptides and underlying SNPs were confirmed by Sanger sequencing. **(B)** Comparison of matched ions observed in candidate spectra (top) and synthetic peptide spectra (bottom). Top 10 most intense ions are labeled, retention time difference and correlation between experimental and synthetic peptide spectrum is reported. *RGP1* peptide is singly charged and was therefore compared to HCD synthetic spectra. **(C)** Base calls and sanger traces of underlying SNPs. Positions of SNPs are indicated by an arrow. *RBBP7* SNP is homozygous.

**Supplementary figure 6:**
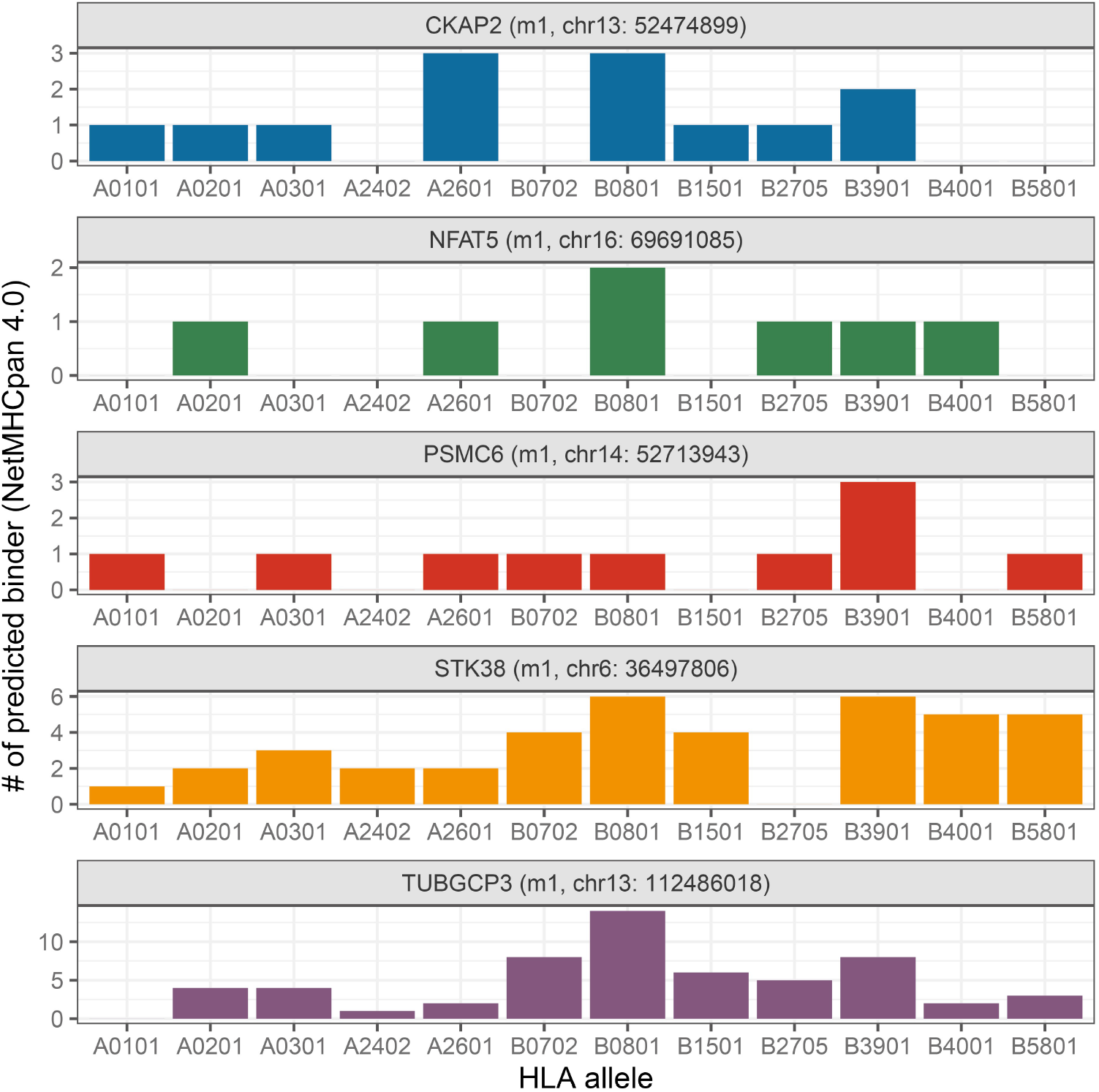
Frameshift parts of source proteins of identified InDel neoepitope generate numerous nonamers predicted to bind HLA supertype alleles. Binding prediction for overlapping nonamers originating from frameshift part of mutated source proteins was performed using NetMHCpan 4.0.

**Supplementary figure 7:**
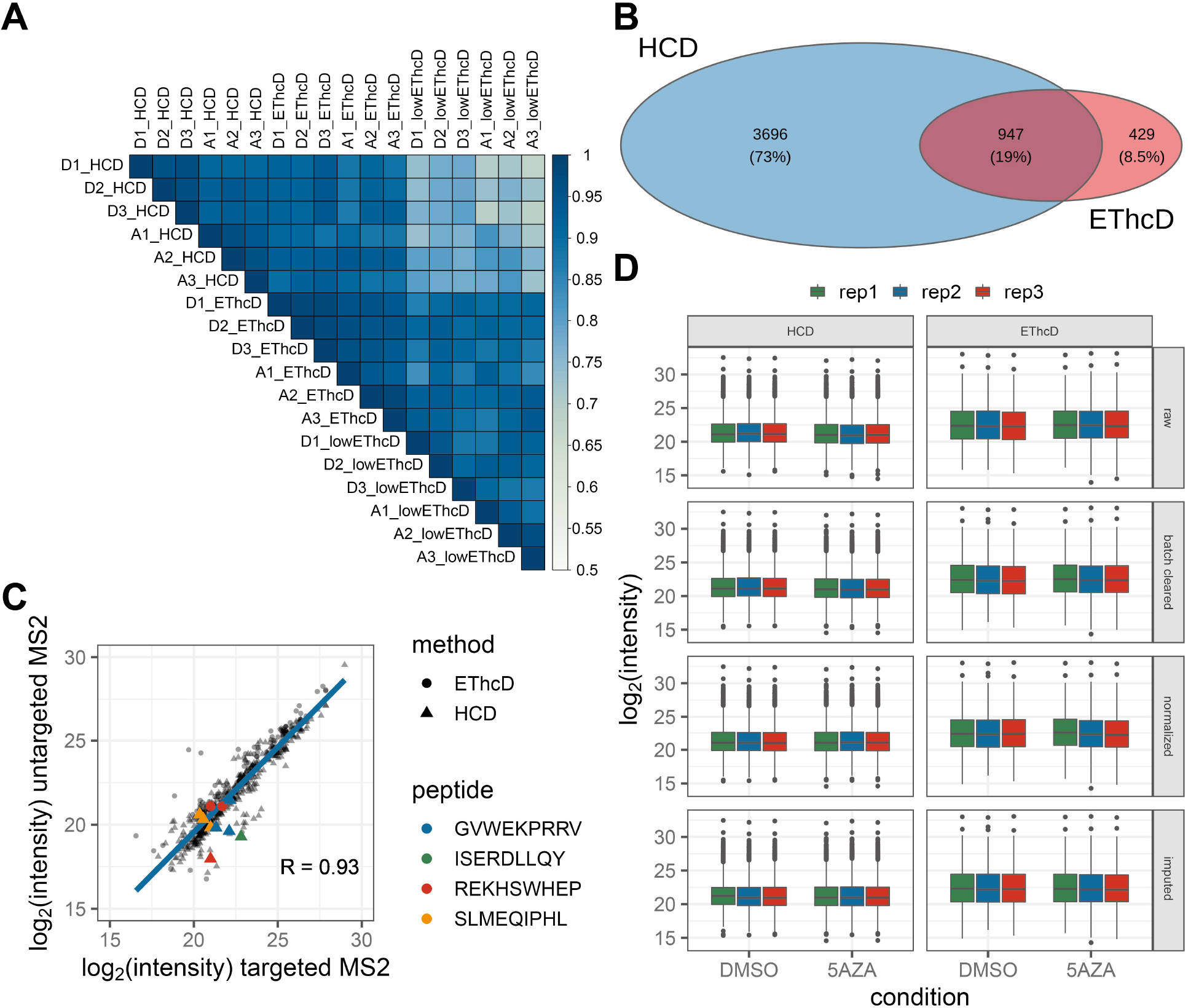
Overview of quantification datasets and dataset processing workflow. **(A)** Correlation matrix showing correlation of raw intensity values between all recorded datasets. Samples are labeled with condition (D = DMSO, A = 5AZA), replicate number (1–3) and MS fragmentation method (HCD, EThcD, lowEThcD). **(B)** Overlap between datasets of quantifiable peptides (measured in at least two out of three replicates per condition) for datasets recorded with HCD and EThcD fragmentation. **(C)** Intensities of peptides identified using a targeted MS2 approach versus intensities of peptides identified using untargeted MS2 approach (HCD/EThcD). Datapoint shape represents MS fragmentation method. Targeted InDel neoepitopes are colored. Peptide KRSSTILRL (*NFAT5*) was not measured with untargeted MS2 approach (HCD/EThcD) **(D)** Data processing overview. Top panel shows distribution of raw intensity values for each replicate, second shows intensities after batch clearing, third after normalization and bottom panel after imputation of missing values.

**Supplementary figure 8:**
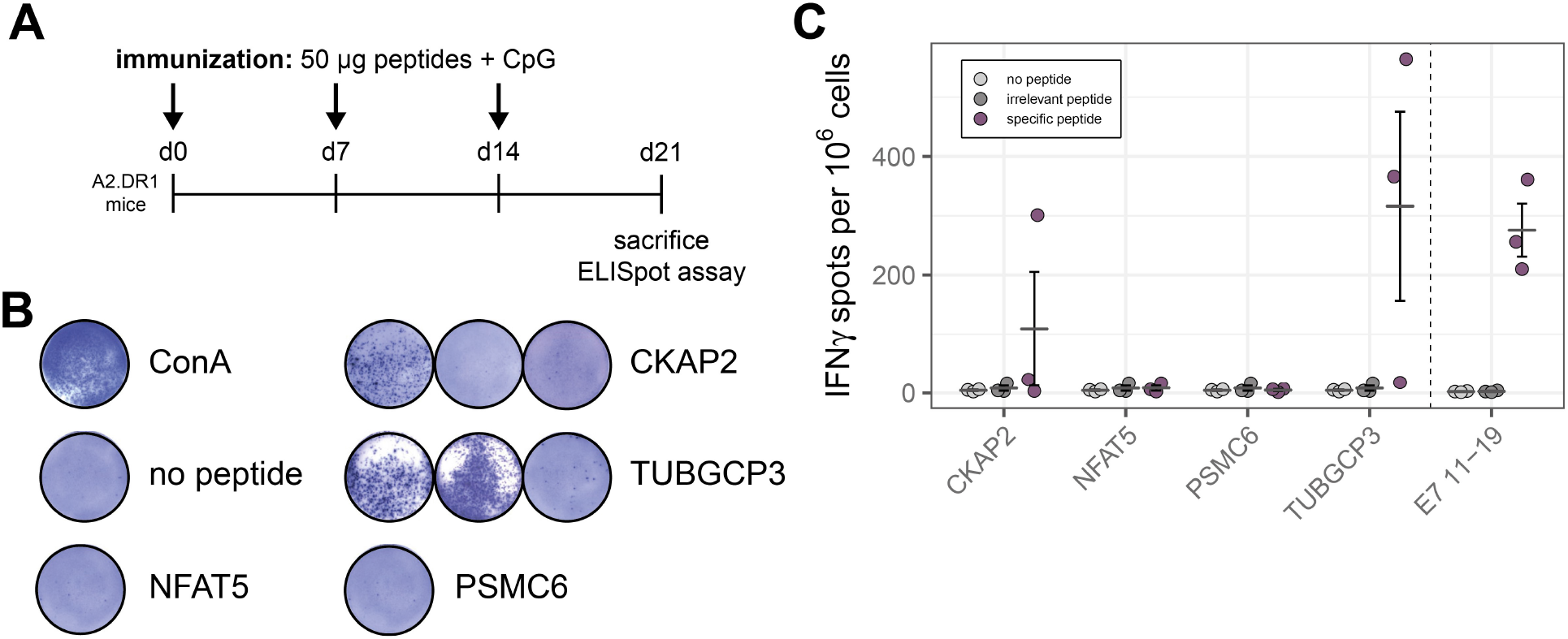
In vivo immunization of A2.DR1 mice with pooled InDel neoepitopes. **(A)** Immunization scheme. **(B)** Representative ELISpot assay results for whole spleenocytes stimulated with ConA (assay positive control), no peptide control, and InDel neoepitopes. **(C)** Quantitative analysis of ELISpot assays. Bars represent mean ± SEM of N = 3.

### Supplementary tables

**Supplementary table 1:** InDel mutation analysis by gel capillary electrophoresis in MSI CRC cell lines. Frequency of underlying frameshift mutations of identified InDel neoepitopes was tested in 24 MSI CRC cell lines. m1, minus one base pair deletion; p1, plus one base pair insertion.

| gene<br>cell line | NFAT5 | TUBGCP3 | PSMC6 | STK38 | CKAP2 |
| --- | --- | --- | --- | --- | --- |
| HCT-116 | m1 | m1 | m1 | m1 | m1 |
| Coga1 | wt | wt | wt | wt | wt |
| Colo60H | wt | wt | wt | wt | wt |
| DLD1 | wt | wt | wt | wt | wt |
| HCT15 | wt | wt | wt | wt | wt |
| HDC135 | wt | wt | wt | wt | wt |
| HDC143 | wt | wt | wt | wt | wt |
| HROC24 | wt | wt | wt | m1 | p1 |
| K073A | wt | wt | wt | wt | wt |
| KM12 | wt | wt | wt | wt | m1 |
| LIM1215 | wt | wt | wt | wt | wt |
| LIM2405 | wt | wt | wt | wt | wt |
| LIM2412 | wt | wt | wt | wt | wt |
| LIM2537 | wt | wt | wt | wt | wt |
| LIM2551 | wt | wt | wt | wt | wt |
| LoVo | wt | m1 | wt | wt | wt |
| LS174T | wt | wt | wt | wt | wt |
| LS411 | wt | wt | wt | wt | m1 |
| RKO | wt | wt | wt | wt | wt |
| TC7 | wt | wt | wt | wt | wt |
| TC71 | wt | wt | wt | wt | wt |
| VaCo457 | wt | wt | wt | wt | wt |
| VaCo5 | wt | wt | wt | wt | wt |
| VaCo6 | wt | wt | wt | wt | m1 |
| total | 1/24 (4.17%) | 2/24 (8.3%) | 1/24 (4.17%) | 2/24 (8.3%) | 5/24 (20.83%) |

**Supplementary table 2:** Target m/z list of InDel neoepitopes for targeted MS2 method.

| peptide | m/z | z | t start (min) | t stop (min) | gene | method |
| --- | --- | --- | --- | --- | --- | --- |
| SLM(ox)EQIPHL | 542.2788 | 2 | 60 | 90 | CKAP2 | HCD, EThcD |
| SLMEQIPHL | 534.2813 | 2 | 60 | 90 | CKAP2 | HCD, EThcD |
| KRSSTILRL | 358.5645 | 3 | 27 | 47 | NFAT5 | HCD, EThcD |
| REKHSWHEP | 302.1507 | 4 | 7 | 27 | PSMC6 | EThcD |
| REKHSWHEP | 402.5319 | 3 | 7 | 27 | PSMC6 | HCD, EThcD |
| ISERDLLQY | 568.8009 | 2 | 60 | 80 | STK38 | HCD, EThcD |
| RLSSCCPVA | 468.2255 | 2 | 41 | 60 | TGFBR2 | HCD, EThcD |
| GVWEKPRRV | 376.2208 | 3 | 15 | 35 | TUBGCP3 | HCD, EThcD |

**Supplementary table 3:**
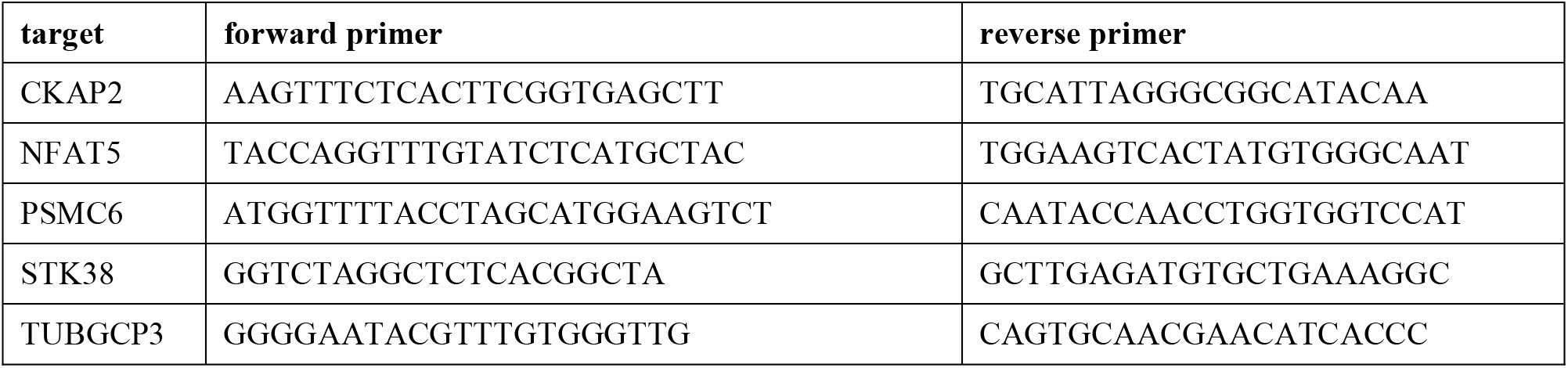
Primer used for validation of underlying frameshift mutations of identified InDel neoepitopes by Sanger sequencing.

**Supplementary table 4:**
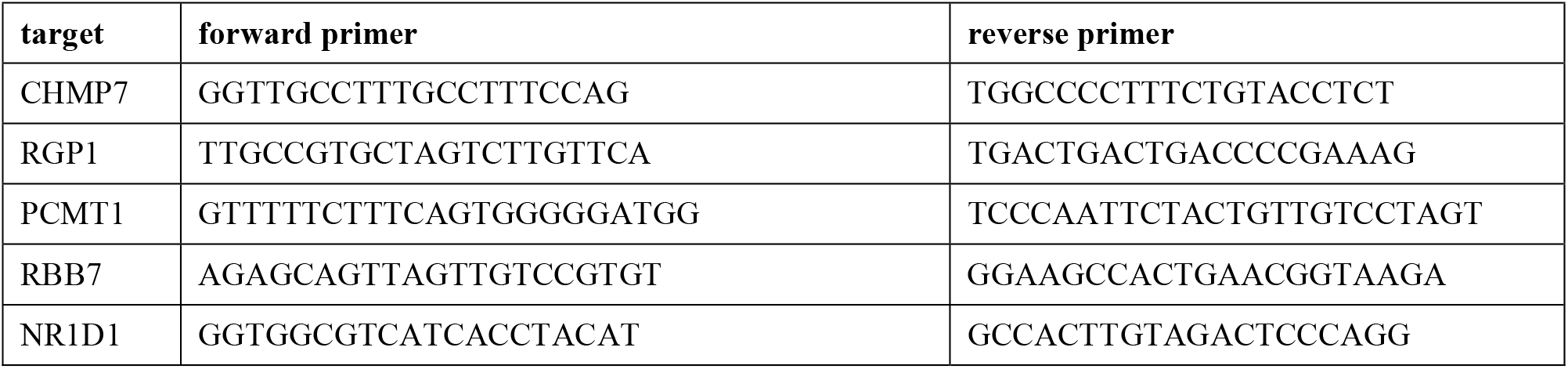
Primer used for validation of underlying SNPs of identified SNP neoepitopes by Sanger sequencing.

**Supplementary table 5:**
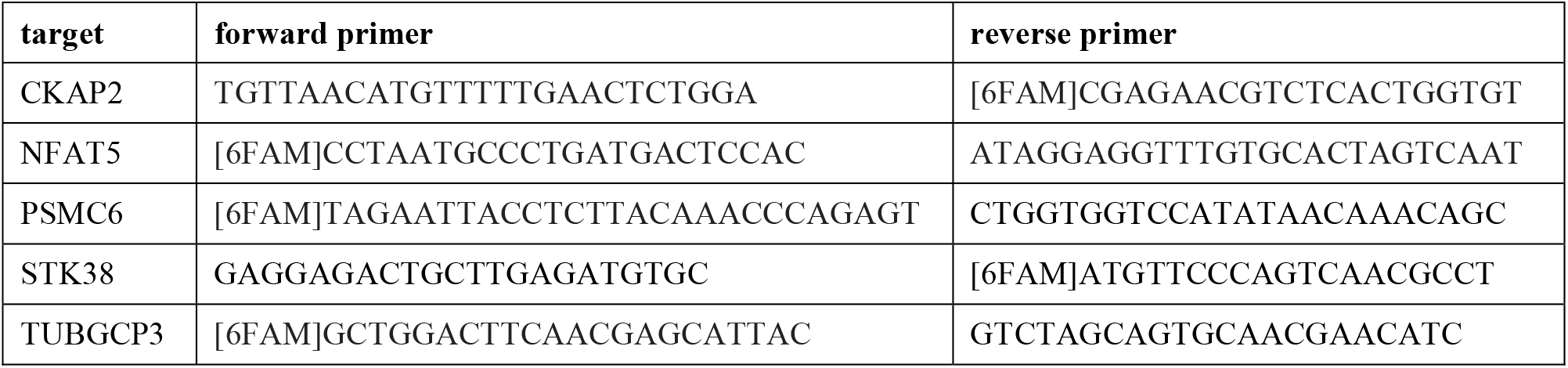
Primers used for mutation analysis of gDNA isolated from MSI CRC cell lines and patients by gel capillary electrophoresis. Labeled primers are indicated.

**Supplementary table 6:**
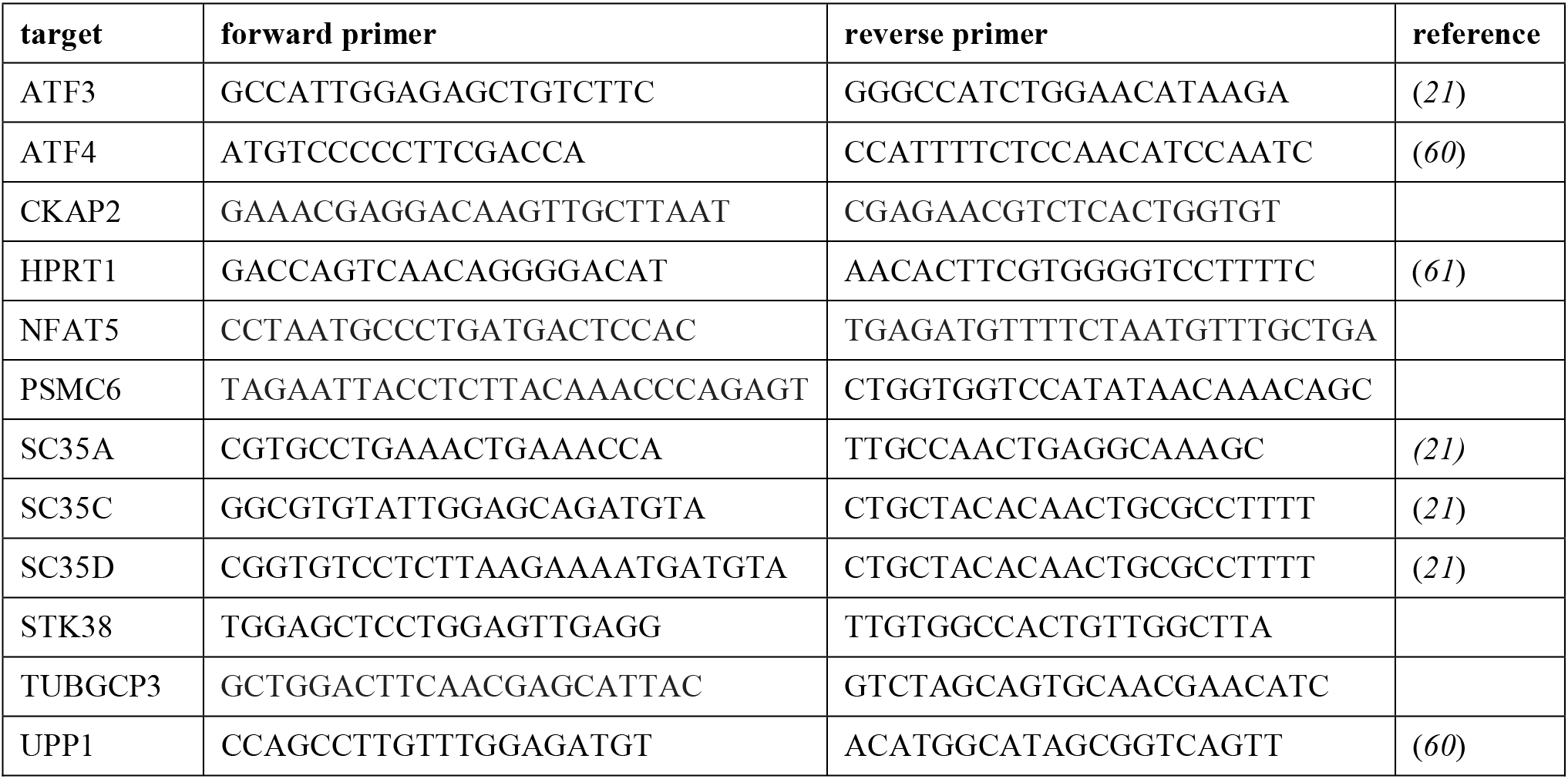
Primer used for qPCR analysis of endogenous NMD targets and frameshift-bearing transcripts.

